# The *Taxu*s genome provides insights into paclitaxel biosynthesis

**DOI:** 10.1101/2021.04.29.441981

**Authors:** Xingyao Xiong, Junbo Gou, Qinggang Liao, Yanlin Li, Qian Zhou, Guiqi Bi, Chong Li, Ran Du, Xiaotong Wang, Tianshu Sun, Lvjun Guo, Haifei Liang, Pengjun Lu, Yaoyao Wu, Zhonghua Zhang, Dae-Kyun Ro, Yi Shang, Sanwen Huang, Jianbin Yan

**Author notes:** Correspondence to (J.Y.); (S.H.). These authors contributed equally to this work (Xingyao Xiong, Junbo Gou, Qinggang Liao, Yanlin Li).

## Abstract

The ancient gymnosperm genus Taxus is the exclusive source of the anticancer drug paclitaxel, yet no reference genome sequences are available for comprehensively elucidating the paclitaxel biosynthesis pathway. We have completed a chromosome-level genome of *Taxus chinensis* var. *mairei* with a total length of 10.23 Gb. Taxus shared an ancestral whole-genome duplication with the coniferophyte lineage and underwent distinct transposon evolution. We discovered a unique physical and functional grouping of *CYP725As* (cytochrome P450) in the Taxus genome for paclitaxel biosynthesis. We also identified a gene cluster in the taxadiene biosynthesis, which was mainly formed by gene duplications. This study will facilitate the elucidation of paclitaxel biosynthesis and unleash the biotechnological potential of Taxus.

**One Sentence Summary:** A chromosome-level genome assembly of *Taxus chinensis* var. *mairei* uncovers its unique genome evolution process and genetic architectures for the paclitaxel biosynthesis pathway.

## Main

Taxaceae (*Taxus spp.*), a widespread nonflowering conifer with substantial economic value, contains six extant genera and over 28 species^1^. *Taxus* is the largest genus in Taxaceae, including common species such as *T. chinensis*, *T. brevifolia,* and *T. baccata*, and it is mainly distributed in Asia, North America, and Europe^2^. For decades, Taxus has served as a natural source to obtain paclitaxel (trade name Taxol), a well-known chemotherapy agent against various cancers^3^. Nevertheless, plant-derived paclitaxel suffers from a short supply due to its low abundance in Taxus, limiting its clinical application. In this instance, multiple strategies have been employed to address supply issues^4^, and promising progress has been made in chemical^5^ and semichemical synthesis^6^, direct extraction from Taxus cell lines^7^, fermentation of endophytic paclitaxel producing fungi^8^, and metabolic engineering of paclitaxel production using heterologous systems^9^.

As a tetracyclic diterpene, paclitaxel is biosynthesized by a complex metabolic pathway^10^. The paclitaxel pathway starts with geranylgeranyl diphosphate (GGPP) synthesis through condensation of isoprenyl diphosphate (IPP) and dimethylallyl diphosphate (DMAPP)^11^. GGPP is then cyclized by taxadiene synthetase (TS), generating a unique diterpene skeleton, taxadiene^12^. Taxadiene is subsequently decorated by a series of reactions including hydroxylation, oxidation, epoxidation, acylation and benzoylation, to generate the final product via catalysis by various enzymes (e.g., hydroxylase, oxidase, epoxidase, oxomutase, and transferase)^13–15^. To date, over twenty enzymes have been identified in the paclitaxel biosynthetic pathway. However, several essential steps such as C1 hydroxylation, C9 oxygenation and oxetane formation in the paclitaxel biosynthetic pathway, remain to be clarified. Moreover, studies have shown that jasmonates (JAs), gibberellin, auxin, and ethylene are involved in the regulation of paclitaxel biosynthesis to maintain a delicate balance between growth and defense in Taxus^16–18^. Several transcription factors (TFs), including AP2/ERF, WRKY, MYC, and MYB, have been found to regulate the expression of paclitaxel biosynthetic genes^19–21^. However, the comprehensive regulatory mechanisms underlying the growth-defense tradeoff are still poorly understood.

Although a complete Taxus genome sequence can provide valuable bioinformatic and genetic resources to understand paclitaxel biosynthesis and regulatory mechanisms in depth, the sizeable and complex genome (2C-value: 22.3-24.3 picogram) of Taxus has hindered its *de novo* draft genome assembly to date^22^. Here, we successfully assembled the Taxus genome, and presented a reference-grade genome sequence of *T. chinensis* var. *mairei* containing 10.23 Gb data with contig N50 of 2.44 Mb, 9.86 Gb of which was assigned to 12 pseudochromosomes. We demonstrated that the CYP725A (cytochrome P450) genes, closely related to paclitaxel biosynthesis, have evolved independently in a unique physical and functional grouping in the Taxus genome. Moreover, we uncovered a gene cluster for taxadiene biosynthesis that contains a new type of taxadiene synthase. These results contribute to our understanding of the biological and evolutionary questions regarding paclitaxel biosynthesis and provide insights into the genome structure and organization of gymnosperms.

## Results

### Taxus genome sequencing, assembly and annotation

To build a chromosome-level genome assembly of Taxus, genomic DNA was extracted from endosperm calli. The endosperm of *T. chinensis* var. *mairei* seeds with haploid chromosomes was used to culture the callus, as it could prevent the influence of heterozygous elements in the genome assembly. *K*-mer analysis showed that the genome size of *T. chinensis* var. *mairei* was approximately 10 Gb (Supplementary Fig. 1), which is consistent with the results from the flow cytometry tests^23^. A *de novo* assembly of the Taxus genome was achieved by PacBio continuous long reads (318.05 Gb) and augmented with Illumina whole-genome sequencing (WGS) reads (693.73 Gb) (Supplementary Table 1). After the application of high-throughput/resolution chromosome conformation capture (Hi-C) (Supplementary Table 2), 9.86 Gb of sequence data could be assigned to 12 pseudochromosomes (Supplementary Fig. 2, Supplementary Table 3), which covered 96.28% of the genome (Supplementary Table 4). We finally obtained the genome sequence with a total length of 10.23 Gb and a contig N50 of 2.44 Mb (Fig. 1a, Supplementary Table 4).

**Fig. 1.**
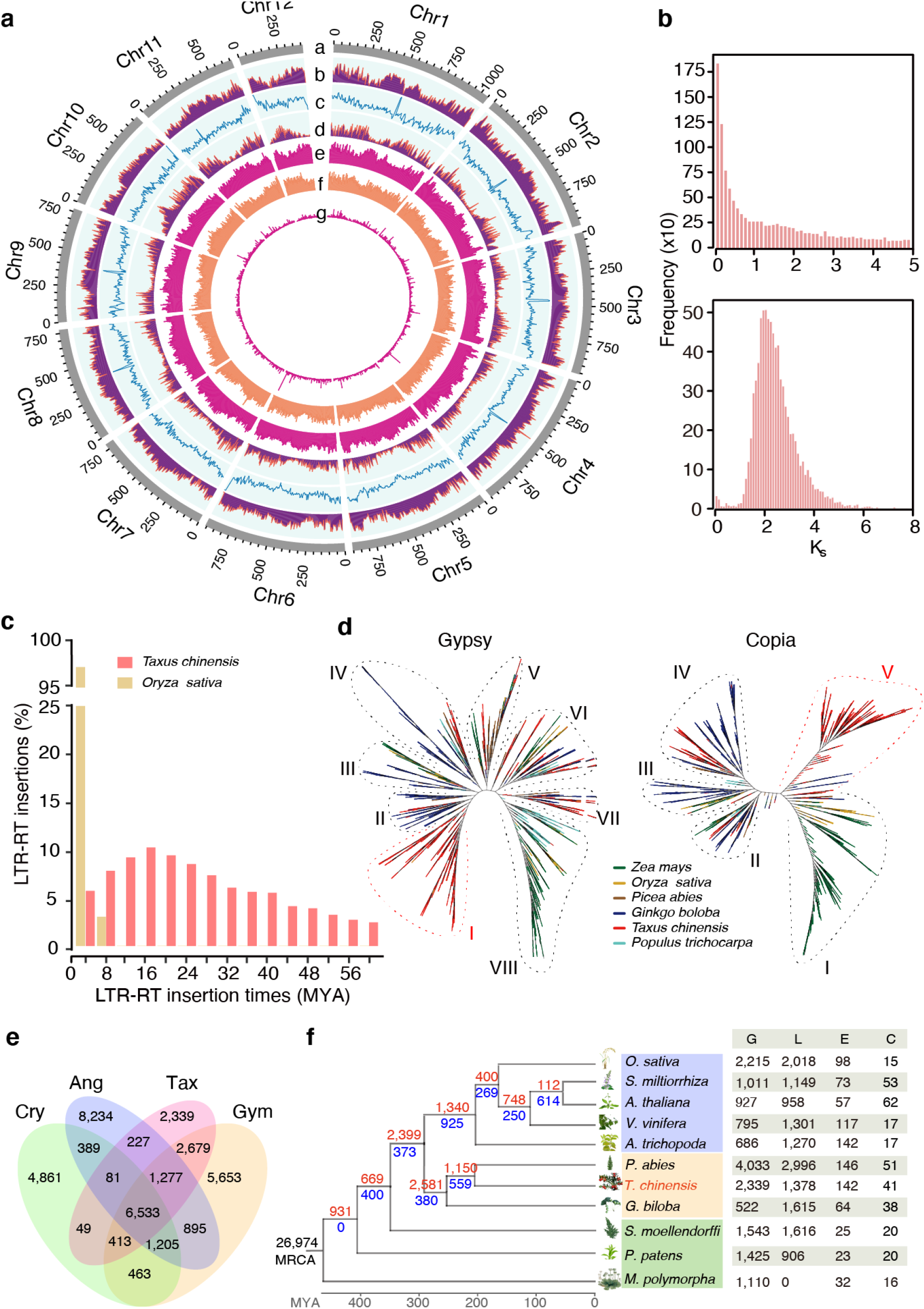
Genomic features of *T. chinensis* var. *mairei*. (a) Genomic landscape of the twelve assembled pseudochromosomes. Track a represents the length of the pseudochromosomes (Mb); b, c, and d represents repeat element density, GC content, and distribution of gene density, respectively; e, f and g show the distribution of Ty3/Gypsy, Ty1/Copia, and unknown LTRs, respectively. These metrics are calculated in 5 Mb windows. (b) Whole genome duplication (WGD) analysis based on the substitution rate distribution of paralogs. Upper panel: Histogram of the *Ks* distribution from Taxus paralogs based on an all-to-all blast to total genes. Lower panel: *Ks* distribution of paralogs based on syntenic analysis. *Ks* values were calculated using the YN model in *KaKs*_calculator. (c) Expansions and diverse sets of LTR elements in the Taxus genome. The histogram shows distributions of insertion times calculated for LTRs in Taxus and rice, using mutation rates (per base year) of 7.3 × 10 ^−10^ for Taxus and 1.8 × 10 ^−8^ for rice. The LTR-RT insertions of *T. chinensis* var. *mairei* and *Z. mays* are shown as columns in different colors. (d) Heuristic maximum like hood trees of Ty3/Gypsy (shown as Gypsy) and Ty1/Copia (shown as Copia) from 6 plant species. The two trees were constructed from amino acid sequence similarities within the reverse transcriptase domains of Gypsy and Copia from 6 plant species. Gypsy elements are divided into eight families (I-VIII), and Copia contains five families (I-V). The representative plants are shown as colored lines. (e) Venn diagram for orthologous protein-coding gene clusters in cryptogam (Cry), angiosperm (Ang), gymnosperm (Gym), and *T. chinensis* var. *mairei* (Tax). The cryptogams include *M. polymorpha*, *P. patens* subsp. *patens*, and *S. moellendorffii*. The angiosperms include *A. trichopoda*, *V. vinifera*, *A. thaliana*, *S. miltiorrhiza*, and *O. sativa*. The gymnosperms include *P. abies* and *G. biloba*. The number in each sector of the diagram represents the total number of genes across the four comparisons. (f) Evolution analysis of gene families in Taxus and selected plants. The red numbers on the branches of the phylogenetic tree indicate the number of expanded gene families, and the blue numbers refer to the number of constricted gene families. The supposed most recent common ancestor (MRCA) contains 26,974 gene families. G, L, E, and C in the table at right represent the number of gains, losses, expansions and constrictions in the gene families among 11 plant species.

**Fig. 2.**
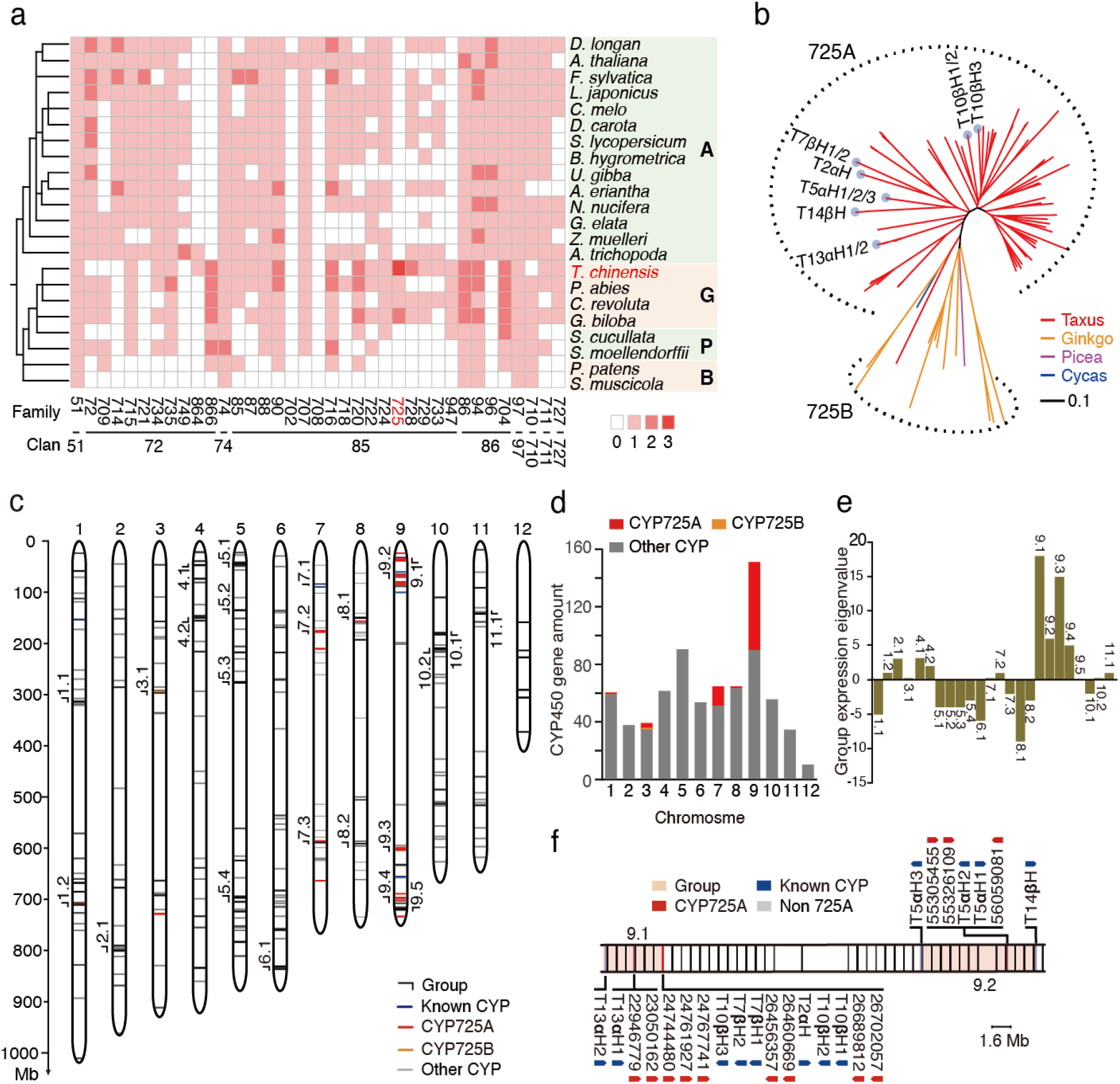
Evolution and genomic architecture of Taxus CYP450s. (a) Phylogenomic analysis of the non-A-type CYP450s in the representative plant species. Abbreviations: A, Angiosperms; G, Gymnosperms; P, Pteridophytes; B, Bryophytes. The color of each block is based on the number of genes in each family, and 0, 1, 2, and 3 indicate that the ranges from 0, 1-10, 10-50, and 50-100 genes, respectively. (b) Phylogenetic analysis of the CYP725 subfamily in *T. chinensis* var. *mairei* (Taxus), *G. biloba* (Ginkgo), *P. abies* (Picea), and *C. revolute* (Cycas). The dotted outline shows the gene spheres of the CYP725A and CYP725B subfamilies. The dusty blue dots on the end of the phylogenetic branches represent the known paclitaxel pathway CYP725A genes and their homologs. The neighbor-joining tree was constructed by Interactive Tree Of Life (iTOL) software. The evolutionary distances were analyzed by the *p*-distance method, and the branch lengths were scaled by the bar. (c) Distribution of CYP450 genes on the twelve pseudochromosomes in Taxus. Each short line on the pseudochromosomes represents a CYP450 gene. CYP725As, CYP725Bs, and the other CYP450s are marked by red, orange and gray lines, respectively. The known CYP450s in the paclitaxel biosynthesis pathway (known CYP) are shown in blue. The CYP450 groups (≥ 7 CYP450 genes and ≤ 5.26 Mb of gene spacing between two adjacent CYP450s) are labeled outside of the corresponding position on the pseudochromosomes. (d) Histogram plot of the number of CYP450 genes on each pseudochromosome. The CYP725 genes (shown in red and orange) were mainly distributed on pseudochromosome 9, while the other CYPs (shown in grey) were distributed randomly on 12 pseudochromosomes. The Y-axis represents the number of CYP450 genes. (e) Group-based gene expression profiles in response to methyl jasmonate (MeJA) treatment. RNA sequencing analysis was performed with the low paclitaxel-yielding cell line (LC) treated with 100 μM MeJA for 4 h. The expression of the gene group was calculated by the sum of the expression levels of each CYP450, and each upregulated and downregulated CYP450 was calculated as 1 and −1, respectively, based on their RPKM (reads per kilobase per million reads) values. (f) Map of CYP725As located in groups 9.1 and 9.2. The ranges of the gene groups on pseudochromosome 9 is marked in pink. CYP725As and the other genes are marked by red and gray vertical lines, respectively. The known CYP450s in the paclitaxel biosynthesis pathway (known CYP) are shown in blue. The arrows show gene orientations.

Based on the genomic information, 42,746 protein-coding genes were further annotated by integrating transcriptome data, homologous alignments, and *ab initio* gene models. 73.02% of the genes (31,214 out of 42,746) could be supported by RNA-seq data (Supplementary Fig. 3, Supplementary Table 5). The BUSCO analysis further demonstrated that 1,052 of the 1,614 core genes were complete, showing relatively high completeness of the assembled genome in gymnosperms (Supplementary Table 6). Furthermore, 36,518 coding genes, accounting for 85.43% of the total predicted genes, were assigned to functional categories with an *E*-value less than 10^−5^ (Supplementary Table 7).

**Fig. 3.**
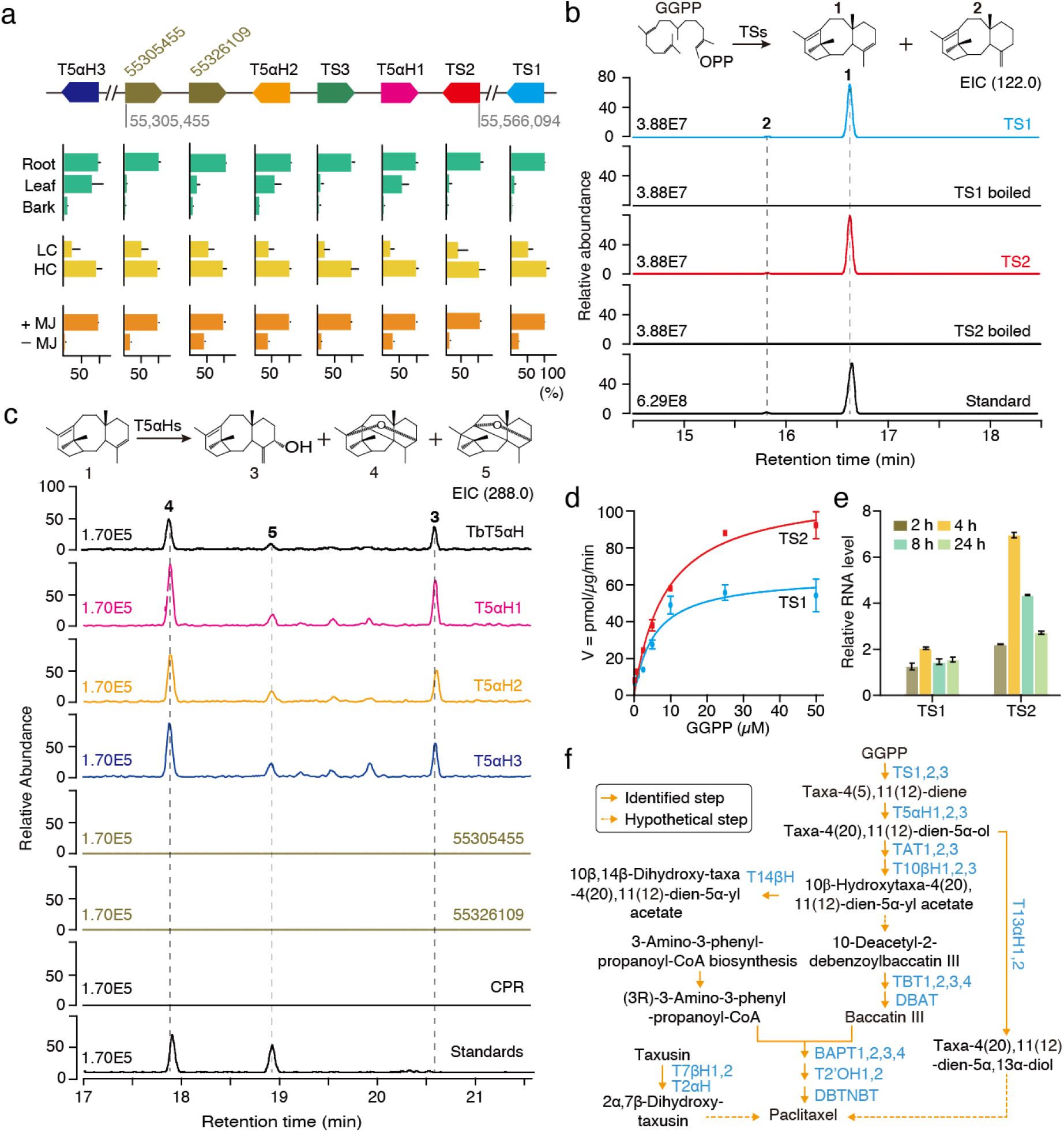
Functional identification of the paclitaxel biosynthesis gene cluster. (a) Genomic architecture and expression pattern of the taxadiene cluster. The arrows indicate the relative position and direction of the genes in the cluster. Here, 55,305,455 and 55,566,094 indicate the starting and ending positions of the cluster on pseudochromosome 9. The two unknown CYP725A genes are represented by their gene starting position 55326109 and 55305455, respectively. TS1 and T5*α*H3 are located at 72,105,619-72,109,598 and 49,866,845-49,868,629 bp on chromosome 9, respectively. The relative expression levels of taxadiene cluster genes in Taxus are based on their RPKM values. The expression level of genes with high sequence similarity was distinguished based on sequencing read counts of the exons that include different bases, and adjusting the alignment threshold to no mismatch. RNA-Seq datasets are from roots, leaves, and bark of male plants (shown in green); two *T. chinensis* var. *mairei* half-sib cell lines, a high paclitaxel-yielding cell line (HC) and a low paclitaxel-yielding line (LC), respectively (shown in yellow); and MeJA-treated LC (+MJ) and MeJA-untreated LC (-MJ) (shown in orange). Data are means ± S.D. (n = 3). (b) Analysis of TS activity *in vitro*. The purified recombinant TS1-His and TS2-His were incubated with the substrate GGPP overnight at 32°C. The reaction products were analyzed by GC/MS. TS catalyzes GGPP to produce a major product (1, taxa-4(5),11(12)-diene) and a minor product (2, taxa-4(20),11(12)-diene), while boiled TSs have no TS activity. *m/z* 122 is a characteristic ion of taxadienes. The taxadiene confirmed by NMR analysis was used as a reference standard (Standard). (c) Analysis of the activity of T5*α*H and two novel CYP725As *in vitro.* The *in vitro* enzyme assay was carried out with the purified taxadiene substrate and yeast microsomes, each including one of the six CYPs (T5*α*H1, T5*α*H2, T5*α*H3, TbT5*α*H, 55326109 or 55305455) and cytochrome P450 reductase (CPR). T5*α*H1/2/3 can produce three oxygenated taxadiene products (3, 5(12)-oxa-3(11)-cyclotaxane (OCT); 4, 5(11)-oxa-3(11)-cyclotaxane (iso-OCT), and **5,** taxa-4(20),11(12)-dien-5*α*-ol), whereas no catalytic compounds were observed for 55326109, 55305455 and CPR. *T. brevifolia* taxadiene 5-*α*-hydroxylase (TbT5*α*H), shown to have taxadiene 5-*α*-hydroxylase activity, was used as a positive control. (d) Kinetic evaluation of GGPP oxidation catalyzed by TS1 (blue circle) and TS2 (red rectangle). *K*_m_ = 5.5 ± 1.6 μM (TS1), *K*_m_ = 8.6 ± 1.5 μM (TS2), *k*_cat_ = 0.0029 s^−1^ (TS1), and *k*_cat_ = 0.0090 s^−1^ (TS2). (e) Quantitative real-time PCR analysis of the transcription levels of TS1 and TS2 in the Taxus cell line LC treated with 100 μM MeJA for the indicated times. The relative gene expression levels are represented as the average fold change (2^*−ΔΔCt*^). The Taxus actin 1 gene (*7G702435613*) was used as an internal reference. Data are means ± S.D. (n = 3). (f) Biosynthesis pathway of paclitaxel in *T. chinensis* var. *mairei*. The solid arrows indicate the identified steps in the paclitaxel pathway, whereas the dotted arrows show the hypothetical steps. TS, taxadiene synthase; T5*α*H, taxadiene 5-*α*-hydroxylase; T13*α*H, taxane 13-*α*-hydroxylase; TAT, taxadien-5-*α*-ol O-acetyltransferase; T10*β*H, taxane 10-*β*-hydroxylase; T14*β*H, taxoid 14-*β*-hydroxylase; T2*α*H, taxoid 2-*α*-hydroxylase; T7*β*H, taxoid 7-*β*-hydroxylase; TBT, 2-*α*-hydroxytaxane 2-O-benzoyltransferase; DBAT, 10-deacetylbaccatinIII 10-O-acetyltransferase; BAPT, baccatin III amino phenylpropanoyl-13-O-transferase; and DBTNBT, 3’-N-debenzoyl-2’-deoxytaxol N-benzoyl transferase.

### Taxus experienced a whole-genome duplication event in the cupressophyte clade

Given that the whole genome duplication (WGD) is a significant evolutionary force contributing to the expansion of plant genome size^24^, we investigated whether Taxus had experienced any WGD events. We built a paralogous gene pair set by performing an all-against-all blastp search. The number of synonymous substitutions per synonymous site (*Ks*) of paralogs was calculated using the gene pair set. As shown in Fig. 1b, the number of *Ks* frequency exhibited an apparent decay without a natural distribution with increasing of *Ks* values, which indicated that no recent WGD event occurred in the Taxus genome. Moreover, we noticed that most of the *Ks* values were less than 0.8, indicating genes duplication in combination with saturation and stochasticity effects may obscure WGD^24^. We further used MCScanX to produce 8,148 syntenic gene pairs from the all-against-all blastp data and entered them into the *Ks* and distance-transversion rate at fourfold degenerate sites (4DTv) calculations. The results showed two signature peaks located at 2.1 for *Ks* (Fig. 1b) and 0.7 for 4DTv (Supplementary Fig. 4), suggesting the presence of an ancient WGD in Taxus. Together with previous studies that revealed an ancient WGD event (WGD-ζ) in the common ancestor of angiosperms and gymnosperms^24,25^, all the results mentioned above suggested that Taxus shared the common ancient WGD with other coniferophyte lineages.

### Taxus genome expansion is linked with retrotransposons

Except for the role of WGD in enlarging the Taxus genome size, we noticed that repetitive sequences constituted a significant component of the Taxus genome (Supplementary Table 8). There was a total of 7.79 Gb of repetitive sequences, occupying 76.09% of the entire genome (Supplementary Table 8). Among these repetitive sequences, long terminal repeat (LTR) retrotransposons accounted for the highest proportion at 52.38% (Supplementary Table 8). The insertion time analysis revealed that LTR insertion was a continuous process, and approximately 40% of the insertions occurred 8 to 24 million years ago (MYA) (Fig. 1c). This feature of continuous insertion in the Taxus genome was significantly different from that in the rice genome, where almost 95% of LTR insertions occurred within the last 5 million years^26^. Considering that LTR insertion in Norway spruce and ginkgo mainly occurred 12-24 and 16-24 MYA^23,25^, the continuous insertion of LTRs might be a common phenomenon in gymnosperms.

To further explore the evolution of LTR in Taxus, we analyzed the phylogenies of LTR retrotransposons in a few representative gymnosperm and angiosperm plants. Amino acid sequence similarities within the reverse transcriptase domain of the Ty3/Gypsy retrotransposons (Gypsy) and Ty1/Copia retrotransposons (Copia) were used to construct phylogenetic trees. As shown in Fig. 1d, the Gypsy superfamily members of the gymnosperms ginkgo and picea were distributed in families II-VII, while those of angiosperms mainly belonged to family VIII. In contrast, Taxus Gypsy elements were not only distributed in families II to VIII but also evolved a highly species-specific family (family I), suggesting the expansion of specific Gypsy elements after Taxus speciation. Similarly, the unique expansion phenomenon in Taxus was also observed in the phylogenies of the Copia superfamily (Fig. 1d). Moreover, family V consisted of only Taxus LTRs in the Copia phylogenetic tree displaying a Taxus-specific amplification burst. In addition, Taxus was distributed in family IV, where the gymnosperms ginkgo and picea were located, and families I-III also contained angiosperms, suggesting that Taxus LTRs were placed in a unique position compared to other selected species. These results suggested that the Gypsy and Copia superfamilies of Taxus have undergone a relatively unique evolutionary pattern, especially the specific Gypsy family I and Copia family V.

### Evolution of gene families and elevated secondary metabolism in Taxus

To understand the context of metabolic networks during Taxus evolution, we compared orthologous genes between Taxus and selected gymnosperms, angiosperms and cryptogams (Fig. 1e). In the 35,298 identified orthologous gene families (Supplementary Table 9), we found that 6,533 gene families were shared by the selected species, illustrating their evolutionary conservation (Fig. 1e). Compared with the selected species, 2,339 gene families were exclusive to Taxus (Fig. 1e). Besides, 1,378 gene families experienced loss, while 142 and 41 families underwent expansion and contraction in the Taxus, respectively (Fig. 1f).

Taxus contains 9,747 unique genes (Fig. 1e, Supplementary Table 10), many of which are enriched in the biosynthesis of specialized metabolites, including terpenes, phenylpropanoids, and flavones (Supplementary Table 11). For instance, 57 gene families were annotated to be cytochrome P450 (CYP450) gene families (Supplementary Table 10). Gene expansion analysis demonstrated that 979 genes were enriched in ADP binding, oxidoreductase activity, flavin adenine dinucleotide binding, transferase activity, and signal transduction, etc. (Supplementary Fig. 5, Supplementary Table 12), with eight gene families being associated with CYP450 (Supplementary Table 13). Pfam functional analysis further showed that the Taxus genes were enriched in CYP450 gene families (PF00067.22, p < 0.01) and TFs (PF13837.6, p < 0.01 and PF00847.20, p < 0.01) (Supplementary Table 14). KEGG analysis indicated that the gained and expanded gene families were enriched in a total of 36 and 41 KEGG pathways, respectively, including one phenylpropanoid (ko00940) and three terpenoid metabolic pathways (ko00900, ko00130, ko00902) (Supplementary Tables 11 and 15).

### Evolution and genomic organization of Taxus CYP450s

Given that CYP450s participate in almost half of the enzymatic reactions in paclitaxel biosynthesis^27^, we analyzed Taxus CYP450 families and identified 649 CYP450 genes from the present genome using the reported HMM model (PF00067). These CYP450s can be divided into two catalogs (A-type and non-A-type). The A-type CYP450s included only the CYP71 clan, which consisted of 17 families and 325 genes (Supplementary Fig. 6 and Supplementary Table 16), while the non-A-type CYP450s contained 12 clans that were composed of 27 families and 324 genes (Supplementary Fig. 7 and Supplementary Table 16). Phylogenomic analyses showed that the CYP750 and CYP725 families were significantly expanded in Taxus compared to 68 other representative species, which covered Zygnematophyceae and Sapindaceae (Fig. 2a, Supplementary Figs. 8 and 9, and Supplementary Table 17). The CYP750 family was reported to participate in the biosynthesis of thujone monoterpene, which is involved in defense responses (e.g., resistance against herbivore feeding)^28^, while CYP725 genes were known to contribute to paclitaxel biosynthesis^29^. Phylogenetic analysis of these CYP725 genes further showed that they could be categorized into the CYP725A and CYP725B subfamilies (Fig. 2b). The CYP725A subfamily (a total of 80 genes) exhibited specificity to Taxus, whereas the CYP725B subfamily was universal in gymnosperm plants (including Picea, Cycas, Ginkgo, and Taxus) (Fig. 2b), which suggested that CYP725A underwent independent evolution in Taxus. Considering that all the previously defined CYP450 genes in the paclitaxel pathway belong to the CYP725A subfamily, these results suggested that the expansion of the CYP725A subfamily played vital roles in the evolution of paclitaxel biosynthesis in Taxus.

We noticed that most CYP725A genes (64%) were located on pseudochromosome 9 (Fig. 2c and 2d), exhibiting a distinct aggregated distribution. Gene aggregation analysis further revealed that the Taxus CYP450 genes were not distributed randomly but trended to organize into different gene groups, 24 of which were detected in the genome (Fig. 2c). We found that nearly all these groups, except groups 1.2 and 5.1, contained gene members from no more than three CYP450 families, and ten groups had only one CYP450 family (Supplementary Table 18), suggesting that the grouping of CYP450 genes on the genome had an obvious family aggregation pattern. Furthermore, as an essential phytohormone in the biosynthesis of secondary metabolites^30^, jasmonate is closely related to the expression regulation of CYP450 genes in Taxus (Fig. 2e). Under jasmonate treatment, 8 groups showed an increased expression level, 12 groups showed the inhibition of gene expression (Fig. 2e, Supplementary Fig. 10 and Supplementary Table 18). These results suggested a tendency that the CYP450s in the majority of groups were coexpressed under jasmonate treatment, implying that the grouping of CYP450 genes had some coordination of physiological functions.

The gene expression levels of four gene groups (group 9.1-9.4) on pseudochromosome 9 were upregulated most significantly in the presence of jasmonate (Fig. 2e). More interestingly, groups 9.1 and 9.2 contained all known CYP725A subfamily genes related to paclitaxel biosynthesis and 12 undefined CYP725As (Fig. 2f). The expression profiles of these two groups of CYP725A genes showed that 88% of CYP725As were highly expressed in roots, 79% of CYP725As were highly expressed in the high paclitaxel-yielding cell line (HC), and 88% of CYP725As were upregulated after jasmonate treatment (Supplementary Table 19), which is consistent with the results on the increased level of baccatin III and paclitaxel (by 3-5 times) in the Taxus cell line under jasmonate treatment (Supplementary Fig. 11). These results suggested that the two groups were likely to contain most of the paclitaxel pathway genes that arose during Taxus evolution.

### Taxadiene biosynthetic genes are arranged in gene clusters

PlantiSMASH^31^ analysis further showed that a gene cluster related to terpene biosynthesis was presented in group 9.2 (Fig. 3a and Supplementary Table 20). The gene cluster contained two TS genes (*TS2* and *TS3*, sharing 99.96% nucleotide sequence identity), two T5*α*H genes (*T5αH1* and *T5αH2*, sharing 98.67% nucleotide sequence identity), and two novel CYP725As (Fig. 3a, Supplementary Table 21, and Supplementary Figs. 12 and 13). Moreover, the genes in the cluster showed a highly coordinated tissue expression pattern and expression consistency in response to jasmonate treatment (Fig. 3a), suggesting that the genes could be functionally related. *TS2* and *TS3* were located adjacent to *T5αH1* and *T5αH2* (Fig. 3a), suggesting that the genes involved in the first two paclitaxel biosynthetic steps were organized by a tandem gene duplication event during Taxus genomic evolution. The *Ks* value of these duplicated genes suggested that this TS-T5*α*H duplication occurred at approximately 1.15 MYA. In addition to the *TS* and *T5αH* genes assembled in the cluster, additional *TS* (*TS1*) and *T5αH* (*T5αH3*) genes are located downstream and upstream of the cluster, respectively (Fig. 3a, Supplementary Figs. 12 and 13). Biochemical assays further confirmed that *TS1/2* and *T5αH1/2/3* have taxadiene synthase activity (Fig. 3b, Supplementary Figs. 14) and taxa-4(5),11(12)-diene-5*α*-hydroxylase activity (Fig. 3c, Supplementary Figs. 14 and), respectively, demonstrating that the copied genes possessed the corresponding enzyme activities in *T. chinensis* var. *mairei*.

We further studied the kinetic properties of TS1 and TS2. The *K*_m_ value of TS2 was approximately 1.5 times higher than that of TS1, but the turnover number (*k*_cat_) of TS2 was approximately 3 times greater than that of TS1, indicating that TS2 has a higher catalytic efficiency than TS1 (Fig. 3d). Moreover, exogenous jasmonate treatment resulted in an obviously higher level of TS2 than TS1 transcripts (Fig. 3e), implying that TS2 could play a role in paclitaxel biosynthesis in response to different environmental and developmental cues via JA signaling. Sequence identity analysis showed that TS2 shared only 77-78% protein sequence identity with TS1 and *T. brevifolia* TS (TbTS), which is much lower than the sequence similarity (over 90%) within the previously reported TS genes (Supplementary Table 21), suggesting that TS2 is a unique TS gene diverged from TS1 and TbTS. Phylogenetic tree analysis further confirmed that a *Taxus*-specific gene duplication event at approximately 33.2 MYA resulted in two distinct types of TS genes (Supplementary Fig. 15), demonstrating that taxadiene synthases were encoded by two types of TS genes resulting from gene duplication events in Taxus. Together, these results suggested that the genes involved in the two initial steps of the paclitaxel biosynthesis pathway were arranged in a gene cluster named the “taxadiene gene cluster”. The taxadiene gene cluster might be formed by gene duplications and neofunctionalization in Taxus and somewhat similar to previous studies on operon-like gene clusters in plants^32,33^.

Furthermore, we established a gene-to-gene coregulation network using three rounds of subtraction screening with RNA-Seq datasets. The network could cover all known paclitaxel biosynthetic genes (Supplementary Figs. 16 and 17, Supplementary Tables 22 and 23), indicating its comprehensiveness and high credibility. We identified 17 CYP725A genes, 3 transferases, and 10 TFs with this network, which was strongly associated with known paclitaxel biosynthetic genes (Supplementary Table 24). Real-time quantitative PCR assays confirmed that the expression of certain genes could be induced by jasmonates (Supplementary Fig. 18), implying that their encoded proteins could be investigated as potential enzymes of paclitaxel biosynthesis. Together, these results outlined the biosynthesis pathway of paclitaxel in *T. chinensis* var. *mairei* (Fig. 3f) and provided valuable genetic resources for improving paclitaxel production through genetic breeding and synthetic biology.

## Discussion

The absence of a chromosome-level genome sequence from Taxus has prevented in-depth phylogenomic studies of Taxus. Our study provides an example assembly of a complex genome in trees using various sequencing technologies on DNA from endosperm calli containing haploid chromosomes. Flow cytometry analysis indicated that the nuclear genome (2n) size of the diploid cells of Taxus was approximately 20.80 ~ 24.85 Gb, nearly twice the haploid genome size evaluated by *k*-mer analysis (Supplementary Fig. 1). The vast majority of HiFi sequences from Taxus leaves (diploid) could be mapped to the haploid genome, up to 95%. Moreover, 75.81% of the 228,762,501 SNPs were, 44.97% of the 847,935 InDels were, and 85.64% of the 64,927 structural variants were heterozygous. Taken together with the low heterozygosity (0.02%) of *T. chinensis* var. *mairei*, these results demonstrated that the haploid genome assembly could basically represent its diploid genome, showing the advantages of using endosperm calli for genome assembly.

We found that the complete BUSCOs only increased from 64.7% to 65.2% when the N50 value was increased from 637 kb to 2.44 Mb during the genome assembly. The low BUSCO value might be due to the limitations of the BUSCO reference dataset. The latest dataset version is embryophyta_odb10 (2020-09-10), containing 1,614 genes from single-copy genes of 50 species, including two bryophytes (*Physcomitrella patens* and *Marchantia polymorpha*), one fern (*Selaginella moellendorffii*), and 47 seed plants (all are angiosperms), but has not included any genes of gymnosperms. Consistently, across all of the reported gymnosperm genomes, except for *Gnetum montanum*, the BUSCO values were not higher than 73%, and four of these genomes had values lower than 51% (Supplemental Table 6). The BUSCO value of the Taxus genome was 65.2%, similar to that of *Ginkgo biloba* (69.4%) and *Pseudotsuga menziesii* (67.8%). To assess the Taxus genome quality more comprehensively, we mapped the Illumina DNA sequencing data (~ 693 Gb) for the genome survey onto the assembled genome and found that up to 99.60% of the sequencing data could be mapped, indicating the integrity of the genome assembly. Moreover, we collected transcriptome data from Taxus organs, comprehensively covering eight tissues and cell lines (root, stem, leaf, bark, male strobili, female strobilus, high-yield paclitaxel cell lines, and low-yield paclitaxel cell lines), and mapped the sequencing data to the Taxus genome. The results showed that the average overall mapping rate of transcriptome data to the genome reached 90.45% (Supplementary Table 25), suggesting the integrity of functional genes in the genome.

The Taxus genome contains 4.08 Gb LTR retrotransposons, including 87.28% Gypsy and 12.35% Copia retrotransposons, and a small proportion of unknown LTRs (0.37%) (Supplementary Table 8). The LTR distribution analysis showed that LTRs tended to be distributed throughout the entire chromosome (Supplementary Fig. 19). In particular, Copia tends to be enriched at the two ends of the chromosomes, while Gypsy is more enriched at the chromosome ends and central areas. Compared with previous studies in groundnuts^34^, the two genomes exhibited obvious differences in LTR distribution. The LTR retrotransposons of the groundnut genome are mainly distributed in the central region of chromosomes, close to the centromere. This difference may come from the large disparity in genome size and the difference between angiosperms and gymnosperms.

The LTR insertions in the Taxus genome mainly occurred 8 to 24 MYA during the long insertion period (4-60 MYA) (Fig.1c and Supplementary Fig. 20), while the primary insertion time of LTRs in spruce and ginkgo was 12-24 and 16-24 MYA within their insertion span from 4 to 64 MYA^25,35^. These results suggested that the Taxus genome has a similar LTR insertion time trend to that in the spruce and ginkgo genomes. The very long insertion time phenomenon might be related to the evolutionary characteristics of gymnosperms. It is generally accepted that gymnosperms belong to slow-evolving plants. Their morphology is highly conserved, which is supported by the high similarity between extant species and fossil records. Previous studies have shown that angiosperms and gymnosperms differ considerably in their mutation rates of molecular evolution per unit time, with gymnosperm rates being, on average, seven times lower than those of angiosperm species^36^. For this reason, an insertion time longer than 60 million years is common in gymnosperm genomes because of the much lower mutation rate. For example, up to 8.27% of LTRs were inserted into the ginkgo genome over 60 million years, and 13.31% of LTRs inserted into the Spruce genome over 60 million years^25,35^.

In addition to CYP450 enzymes, acetyltransferases also play an essential role in paclitaxel biosynthesis. We found 127 BAHD acyltransferases by identifying their conserved motifs (HXXXD and DFGWG) (Supplementary Fig. 21). The BAHD acyltransferases in the Taxus were mainly distributed in Clades I, II, VI, and V. Clade V can be divided into three groups (Groups I-III), among which Group I contains all known BAHD acyltransferases in the paclitaxel biosynthesis pathway. It would be worthwhile to investigate whether group I contains other acyltransferases that function in the paclitaxel biosynthesis in the future (Supplementary Table 26). PlantiSMASH analysis indicated that the acyltransferase genes are not organized into any gene clusters. Genomic location analysis showed that the BAHD acyltransferase genes in paclitaxel biosynthesis were mainly distributed on chromosomes 1 and 9 (Supplementary Fig. 17). Furthermore, TAT2 was colocalized with CYP450s in gene group 9.2 (Fig. 2c and Supplementary Fig. 17). The relationship between CYP725As and acetyltransferases in paclitaxel evolution is an interesting aspect to study in the future.

In the Taxus genome, a total of 34 gene clusters related to secondary metabolism were found, including 13 saccharides, seven terpenes, one alkaloid, one saccharide-terpene, one saccharide-polyketide, one lignan-terpene, one terpene-alkaloid, and nine putative gene clusters (Supplemental Tables 20 and 27). Two gene clusters (clusters I and II) belonging to the terpene cluster were involved in paclitaxel biosynthesis because cluster I contained the TS2, TS3, T5αH2, and T5αH3 genes, and cluster II included TS1. Except for these five genes, other related enzymes in the paclitaxel synthesis pathway were not included in any gene clusters. However, we found that most of the known genes involved in paclitaxel biosynthesis, including TAT2, DBAT, TS1/2/3, T7βH1/2, T13αH1/2, T10βH1/2/3, T5αH1/2/3, and T14βH, are located on a small 71.82 Mb region on chromosome 9 (designated T13αH2-DBAT segment: 19994572-91,811,351 bp, Supplemental Fig. 22). Therefore, many genes that play roles in different steps of the paclitaxel synthesis pathway are located in a limited genomic region, implying that there might be a certain coordinated regulatory mechanism of their gene expression. It would be an important future project to investigate whether the genes are organized in a larger-scale gene cluster to achieve better collaborative expression.

To date, all known TS enzymes are homologous to TS1 (amino acid homology > 90%) (Supplementary Table 21A). Our study demonstrated that taxadiene synthases could be encoded by two distinct types of TSs resulting from gene duplication events in Taxus. As a representative of the new type of TS enzyme, TS2 only has approximately 77-78% amino acid homology with the reported TS enzymes (Supplementary Table 21). It exhibits a higher catalytic efficiency than TS1 (Fig. 3d) and more robust induced expression characteristics in treatment with jasmonates (Fig. 3e). The different properties of these two types of TS enzymes implied a new Taxus defense regulation mechanism. In Taxus, the excessive synthesis of taxanes is not conducive to its growth or development although chemicals play an essential role in defense responses. Therefore, it is necessary to accurately and efficiently control the taxane level in cells in response to environmental changes. Our results provide a new hypothesis to explain the regulation of taxane levels in plant cells. When there are no biotic or abiotic stresses, jasmonate signaling is blocked, and TS1 is responsible for taxane biosynthesis to maintain taxanes at a basic level. However, once insect attack or other stresses occur, jasmonate signaling will be activated, and TS2 is rapidly expressed to quickly increase the taxane content in cells.

In addition, we tried to explore the application potential of TS2 in bioengineering. Bian et al. reported an engineered *E.coli* strain with TbTS (belonging to Type I) for the taxadiene product^37^. We replaced the TbTS gene with the TS2 gene (Supplementary Fig. 23). After 60 hours of fermentation, we were surprised to find that the taxadiene titer from the strain containing TS2 was over ten times higher than that containing TbTS, while the OD_600_ of the two strains was not much different (Supplementary Fig. 23). The result showed the great potential of TS2 in bioengineering to produce taxadiene in the future.

We also explored the function of two unknown CYP725As (55305455 and 55326109) in the taxadiene cluster using the well-established T5αH reaction assay. We incubated yeast microsomes that included 55305455, T5αH1, and CPR with the taxadiene as substrate at the same time and analyzed the reaction products by GC-MS. As shown in Supplemental Figure 24, we only detected OCT, iso-OCT, and taxa-4(20),11(12)-dien-5α-ol, which can be obtained by catalyzing taxadiene by the T5αH1 enzyme. The same result was obtained with 55326109 protein in the reaction system. These results suggested that the unknown CYP725s are not involved in the subsequent reaction catalyzed by T5αH. However, the tissue expression specificity of 55305455 and 55326109 was similar to that of TS and T5αH in the cluster, and both of them exhibited higher expression levels in roots than in leaf and bark tissues (Fig. 3a). The real-time PCR assay validated that their expression was induced by jasmonate in Taxus cells (Supplementary Fig. 18), which is consistent with paclitaxel accumulation (Supplemental Fig. 11). Moreover, the updated gene-to-gene coregulation network showed that 55305455 and 55326109 were correlated with DBAT and T5αH1, respectively (Supplemental Fig. 16). These results indicated that the two CYPs may play a role in paclitaxel biosynthesis and metabolism, and are worthy of in-depth study in the future.

## Methods

### Plant materials and genome sequencing

Seeds of a single female *T. chinensis* var. *mairei* were collected from the natural range of Taxus (GPS: N 113° 89’ 55”, E 28° 26’ 32”) in the Liuyang region, Changsha city, Hunan Province, China, in November 2015. Single embryos and endosperm were induced as calli^23,38^.

For sequencing of the haploid tissue, DNA was extracted from the endosperm callus of *T. chinensis* var. *mairei^23^*. The DNA quality was checked by agarose gel electrophoresis and a Qubit fluorimeter (Thermo Fisher, Inc.). The paired-end libraries with a 500 bp insert length were prepared by following the Illumina protocols. Sequencing of the library was performed on the Illumina HiSeq 2500 system. For the PacBio Sequel analysis, SMRTbell TM libraries were prepared according to the manufacturer’s protocol of the sequencing platform. Four independent Hi-C libraries were constructed and sequenced on an Illumina HiSeq 2500 (PE125 bp) at Annoroad Gene Technology Co., Ltd. (Beijing, China).

For circular consensus sequencing (CCS), genomic DNA was isolated from plant leaves using the ctyltriethyl-ammnonium bromide method. A 15 kb DNA SMRTbell library was constructed and sequenced on a Pacbio Sequel II platform, the sequencing reads known as highly accurate long reads, or HiFi reads.

### Genome assembly and gene annotation

The uncorrected PacBio reads were assembled using wtdbg2^39^, the fastest sequence assembler for long noisy reads. The assembly reached the best continuity with the following parameters: -k 0 -p 19 -K 5000 -S1 --aln-noskip --tidy-reads 5000 --edge-min 2 --rescue-low-cov-edges. The software Arrow in the GenomicConsensus package (https://github.com/PacificBiosciences/GenomicConsensus) was applied to generate the consensus sequences from the primary assembly. The raw PacBio reads were aligned to the assembly of red bean^39^ using pbalign (v0.3.1) with default parameters, and then the alignment was passed to Arrow (v2.2.2) to produce the corrected assembly. The consensus process was performed iteratively twice. Further polishing of the assembly genome was conducted using Pilon^40^ with Illumina data, with the following parameters: --fix all --mindepth 0.4 --K 65 --threads 24 --minmq 30 --minqual 30 -- changes.

For Hi-C assembly, the clean Hi-C sequencing data were mapped to the genome draft by HiC-Pro (V2.7.8)^41^, and the library quality was assessed by counting the number of uniquely valid paired-end reads. Only unique valid paired-end reads were maintained for downstream analysis. We used Hi-C data to align and correct the contigs for misassembly through Juicer^42^ pipeline and 3D-DNA pipeline^43^. The assembly package, Lachesis^44^ was applied to perform clustering, ordering and orienting based on the normalized Hi-C interactions. For each pseudochromosome group, the exact contig order and directions were obtained through a weighted directed acyclic graph (WDAG). We filled the gaps among contigs in the pseudochromosomes using TGS-Gapcloser (v1.01)^45^ by two rounds with CLR and HiFi data (26 Gb), respectively. After the filling progress, we further removed the redundant contigs that were not anchored to the chromosomes by Purge Haplotigs (v1.03)^46^.

For assembly assessment, the RNA-seq reads of eight tissues, including mfs, mfl, mfb, mfr, mtf, mtl, mtb and mtr, and HC and LC were mapped to assess the assembly quality (mfs: female strobilus, mfl: female leaf, mfb: female bark of stem, mfr: female root; mtf: male strobili, mtl: male leaf, mtb: male bark of stem, and mtr: male root). The average mapping rate was calculated with the HISAT2^47^ with parameters --score-min L, 0, -0.1.

For repeat annotation and analyses, repetitive elements in the Taxus genome were identified through a combination of *de novo* and homology-based approaches. *De novo* prediction of repeat elements (RE) was carried out using RepeatModeler (v1.0.1, http://www.repeatmasker.org/RepeatModeler/). For homology-based annotation, the RE libraries from Repbase^48^, TIGR (The Institute for Genomic Research)^49^ and the annotated *Ginkgo biloba* genome were merged with the *de novo*-derived library to create the whole dataset. The dataset was then used to mask identified TEs in the Taxus genome with RepeatMasker (v4.0.5, http://www.repeatmasker.org). We identified LTRs by referring LTR_retriever method^50^. Specifically, LTR_finder^50^ and LTRharvest^51^ were first used to identify all the existing LTR sequences in the Taxus genome according to the basic sequence rules of LTRs. The candidate LTR RTs were filtered to remove non-LTR RT repeat elements or those with large amounts of tandem repeats or gaps. Especially in fragmented genome assemblies, these requirements hugely reduce the number of LTR RT candidates but ensure that only full-length LTR RTs are analyzed. We integrated the results and discarded false positives by the LTR_retriever pipeline and then estimated insertion times based on T=D/2μ, where D is the divergence rate and μ is the neutral mutation rate (7.34573×10^−10^)^36^.

For the annotation of protein-coding genes, gene structure prediction was performed using *ab initio,* homology-based and RNA-seq-based pipelines. For the *ab initio* annotation, SNAP^52^, Augustus^53^ and GlimmerHMM were applied. Eight species, *Arabidopsis thaliana*^54^, *Oryza sativa*^55^*, Gnetum montanum*^56^, *Picea abies*^25^*, Ginkgo biloba*^35^*, Selaginella moellendorffii*^57^*, Pinus taeda*^58^ and *Amborella trichopoda*^59^ were chosen for homology annotation to predict protein-coding genes by GeneWise^60^. To generate annotation results based on transcripts, RNA-seq alignment files were generated using TopHat2^61^ and assembled by Cufflinks^62^, and the program PASA^63^ was used to align spliced transcripts and annotate candidate genes. Finally, gene models predicted from three approaches were merged by EVM^64^. The functions of protein-coding genes were identified by mapping sequences against the Gene Ontology^65^, InterProScan^66^, Swiss-Prot (http://www.uniprot.org/)^67^, EMBL database^68^, and TAIR databases^69^.

### Identification of whole genome duplication

Genome-wide duplications were searched in the Taxus genome. Self-alignment of the assembled genome sequence was performed using metablast as described previously^70^. All-*vs*-all paralog analysis in the Taxus genome was performed using reciprocal best hits from primary protein sequences by self-Blastp in Taxus. Reciprocal best hits are defined as reciprocal best Blastp matches with an *E*-value threshold 1e-5, c-score (Blast score/best Blast score) threshold of 0.3^71^, and alignment length threshold of 100 amid acids. The synonymous substitution rate (*Ks*) of reciprocal best hit gene pairs was calculated based on the YN model in *KaKs*_Calculator v2.0^72^. Synteny analysis was performed on Taxus protein-coding genes using MCScanX^73^ to identify whole genome duplication events with default parameters from the top ten self-blastp hits. The synonymous substitution rate (*Ks*) and 4DTv were calculated for Taxus syntenic block gene pairs.

### Genome mining for CYP450s and gene clusters

For the identification and classification of CYP450 genes, hmmsearch was used to identify CYP450 genes in the Taxus genome with PF00067 from the Pfam database^74^. Classification of the 649 CYP450 genes was executed by alignment with the CYP450 database^75^ using standard sequence similarity cutoffs, with definite standards of 97%, 55% and 40% for allelic, subfamily and family variants, respectively. According to the standardized CYP450 nomenclature^76^, CYP450s were divided into A-type and non-A-type CYP450s, and phylogenetic analysis of CYP450 genes was performed for A-type and non-A-type CYP450s. Neighbor-joining phylogenetic trees were constructed using the MEGA7 package with homologous amino acid sequences^77^.

For genome mining for gene clusters involved in plant specialized metabolism, PlantiSMASH^31^ was used to search for potential gene clusters using the default parameters and the GFF (General Feature Format) annotation files of the software. Gene groups were identified by in silico analysis based on the following criteria: 1) the distance between two adjacent CYP450 genes in one group should be less than 5.26 Mb; and 2) one group should contain at least 7 CYP450 genes.

### RNA-seq data analysis for candidate genes in the paclitaxel biosynthesis pathway

All tissues, including female strobilus (mfs), leaf (mfl), bark of stem (mfb), and root (mfr) and male strobili (mtf), leaf (mtl), bark of stem (mtb), and root (mtr), were mapped to the Taxus genome, and the FPKM value was calculated using HISAT2 and StringTie^76^. Expression data from female bark, female roots and female leaves were used to identify the genes associated with paclitaxel biosynthesis. First, we selected genes that were highly expressed in roots or bark compared to leaves. Second, the genes were further confirmed in two Taxus half-sib cell lines (HC and LC) with distinct accumulation patterns of paclitaxel. The genes should be significantly highly expressed in HC. The differentially expressed genes were filtered using edgeR^77^ with logFC >1 and FDR < 0.05. Finally, we obtained 1,638 genes that met the thresholds above. Gene-to-gene networks were constructed using the expression matrix from MeJA-induced cell line (0, 2, 4, 8, 24 h) RNA-seq data. Pearce correlation analysis was performed with the known functional genes as the target genes. Hypothesis development for the Pearson correlation was performed, and pairs with a P-value < 0.05 remained.

### Functional characterization of TS genes

The ORFs (open reading frames) of *TS1* (*ctg6088_gene.1*) and *TS2* (*ctg5306_gene.4*) were cloned by reverse transcription PCR (RT-PCR) from the HC cell line. Plant-Ploc (http://www.csbio.sjtu.edu.cn/bioinf/plant-multi/), ChloroP (http://www.cbs.dtu.dk/services/ChloroP/), and TargetP (http://www.cbs.dtu.dk/services/TargetP/) were used for the prediction of the plastidial target sequence. The 60-residue N-terminally truncated *TS1* and *TS2* genes were inserted into the *Escherichia coli* (*E. coli*) expression vector pET28b to form the constructs pET28b::*TS1* and pET28b::*TS2*, respectively. All expression plasmids were constructed using the Hieff Clone^TM^ One Step Cloning Kit (YEASEN, China) and the primers used in this work are given in Supplementary Table 28. For the *in vitro* enzyme assay, the enzyme assays were performed in a final volume of 500 μL buffer (25 mM HEPES, pH 8.5, 10% glycerol, 5 mM DTT, 5 mM sodium ascorbate, 5 mM sodium metabisulfite and 1 mM MgCl_2_) containing 100 μg of purified protein and 100 μM GGPP (Sigma-Aldrich). The reaction mixture was overlaid with 500 μL of pentane (Macklin, GC-MS grade) and incubated overnight at 32°C. In addition, the mixture was vortexed, and the pentane overlay was subsequently removed by centrifugation at 5,000 rpm for 10 min and concentrated by N_2_ gas before gas chromatography mass spectrometry (GC/MS) analysis. Inactivated TSs-His6 was used as the control. Taxa-4(5),11(12)-diene (**1**) and taxa-4(20),11(12)-diene (**2**) preparations were performed according to a previous study with the taxadiene-producing *E. coli* strain T2 (harboring pMH1, pFZ81, and pXC02)^37^. The organic solutions containing crude compounds **1** and **2** were concentrated on ice under N_2_ gas and re-dissolved in dimethyl sulfoxide (DMSO) for the purification of compounds **1** and **2** by thin layer chromatography (TLC). The purity and concentration were determined by GC-MS (Supplemental Fig. 25).

For the determination of kinetic parameters, standard enzyme assays were carried out in a total volume of 100 μL containing buffer (25 mM HEPES, pH 8.5, 10% glycerol, 5 mM DTT, 5 mM sodium ascorbate, 5 mM sodium metabisulfite and 1 mM MgCl_2_), 34 μg (TS1) or 18 μg (TS2) of recombinant proteins and seven different concentrations of GGPP (0.2, 0.5, 1, 2.5, 5, 10, 25 and 50 μM), which were spiked with [^1-3^H]-GGPP (American Radiolabeled Chemicals, Inc, 30 Ci mM^−1^). Hot [^1-3^H]-GGPP was diluted 400 times using cool GGPP (Sigma, 1 mg mL^−1^). The reaction mixtures were incubated at 32°C for 30 min and then quenched for 10 min using 100 μL of stop solution (containing 1 M EDTA and 4 M NaOH). The reaction mixture was extracted with 800 μL of n-hexane (vortexed for 10 s and 12,000 rpm for 2 min), and 400 μL of the n-hexane layer was subsequently removed and mixed with 2 mL of the liquid scintillation cocktail. The total radioactivity of the reaction products was measured using a liquid scintillation counter (Tri-Carb 2910TR, Perkin Elmer). The kinetic constant was calculated by a nonlinear regression fit to the Michaelis-Menten equation using OriginPro® 8.6 (OriginLab Co., Northampton)^78^.

*E. coli* TS2 was constructed by replacing *TbTS* with *TS2* based on the previous taxadiene-producing *E. coli* TbTS (harboring pMH1, pFZ81, and pXC02, and coexpressing nine genes *AtoB*, *ERG13*, *tHMG1*, *ERG12*, *ERG8*, *MVD1*, *IdI*, *GGPPS* and *TbTS* in *E. coli*)^37^ (Supplementary Fig. 20a). The *E. coli* strains T2 and TS2 were cultivated in 50-mL flasks containing 30 mL of LB medium at 37°C with 100 mg L ^−1^ ampicillin (AMP), 50 mg L^−1^ kanamycin (KAN), and 34 mg L^−1^ chloramphenicol (CM). When the OD_600_ reached approximately 0.1, 1 mM isopropyl *β*-d-1-thiogalactopyranoside (IPTG) was added to the cultures along with 3 ml of dodecane, and then, the bacteria were cultivated at 28°C. Experiments were repeated four times. For cell concentration (OD_600_) and taxadiene measurement, 100-μL cultures and 30-μL organic layers were collected at set intervals (at 8, 13, 22, 37, 48, 60 and 72 hours). The produced taxadiene was detected with GC-MS and quantified with the nonyl acetate standard (Aladdin).

## Acknowledgments

We particularly thank Tiangang Liu, from Wuhan University, for sharing plasmids that were used for the preparation of taxadiene in *E. coli*. We thank Yansheng Zhang (Shanghai University), and Shifeng Cheng (Agricultural Genomics Institute at Shenzhen) for helpful discussion. We are grateful for valuable help with TS evolution from Tao Feng (Wuhan Botanical Garden, Chinese Academy of Sciences).

## Author contributions

The ideas for the paper were conceived by J.Y., S.H., and X.X.; Q.L., G.B., Q.Z., X.W.,T.S., Z.Z., and Y.W. performed bioinformatic analysis; Y.L., X.X., C.L., D.R., L.G., and P.L. prepared the Taxus sequencing samples and the Taxus cell lines; J.G., H.L., D.R., Y.S. and R.D. undertook enzymatic analysis and mass spectrometry; J.Y., H.S., J.G., Q.L., Y.L. and X.X. interpreted the data and wrote the paper.

## Data and materials availability

The raw data used in this study have been deposited in the Genome Sequence Archive at the BIG Data Center, Beijing Institute of Genomics (BIG), Chinese Academy of Sciences and are accessible at http://bigd.big.ac.cn/gsa under bioproject PRJCA003841.

