## Supplemental information for "The *Taxu*s genome provides insights into paclitaxel biosynthesis"

**This PDF file includes:**

1. Supplementary Methods

2. Supplementary Figures 1-26

3. Supplementary Tables 1-8, 24

4. Supplementary References 1-38

### 1. Supplementary Methods

#### RNA sequencing (RNA-seq) for genome structure annotation and expression analysis

Samples of *T. chinensis* var. *mairei* from one male plant and one female plant collected in Hunan Province (GPS: N 113° 89′ 55″, E 28° 26′ 32″) were used for transcriptome sequencing. Eight tissues (female strobilus (mfs), leaf (mfl), bark of stem (mfb), and root (mfr) from the female plant, and male strobili (mtf), leaf (mtl), bark of stem (mtb), and root (mtr) from the male plant) were collected individually. Two half-sib cell lines (HC, LC) were induced from the embryo of the female plant mentioned above; the taxoid contents of the lines were different from each other[1]. The total RNA of these materials was isolated independently using TRIzol® reagent (Invitrogen). The integrity and purity of total RNA were assessed using horizontal agarose electrophoresis and an Agilent Bioanalyzer 2100 (Agilent Technologies, INC). The NEB Next UlturaTM RAN Library Prep Kit (NEB, USA) was used to construct the sequencing library. All RNA-seq libraries were sequenced using the Illumina 2500 platform. The details of the RNA-seq data of each sample are given in Supplementary Table 26.

#### Genome survey by k-mer analysis

Analysis of *k*-mer frequency was adopted to estimate the genome size by using WGS Illumina reads. The *k*-mer frequency was calculated by Jellyfish (v 2.0.0)[2,] and the genome size was estimated based on the formula (Total *k*-mer counts)/(Peak depth).

#### Long-read variant calling for genome assessment

The HiFi reads were advantageous for variant calling because they can span repetitive or other problematic regions. HiFi reads of the Taxus leaves (diploid) were aligned to the Taxus assembly genome to assess the quality. Calling the variants were used to check the coverage of the genome, we aligned HiFi sequencing reads with Minimap2[3] and called SNP (single nucleotide polymorphism) and InDel with bcftools (v1.9)[4]. Then, the SVs (structure variants) were called using Sniffles (v1.0.12) (https://github.com/petabi/sniffles).

#### Analysis of the insertion time of LTR elements

The candidate LTR-RTs were identified using LTR_Finder (v1.02)[5] (parameters: -D 2000 -d 1000 -L 3500 -l 100 -p 20 -C -M 0.8) and LTRharvest[6] (parameters: -similar 90 -vic 10 -seed 20 -seqids yes -minlenltr 100 -maxlenltr 7000 -mintsd 4 -maxtsd 6 -motif TGCA -motifmis 1). Only full-length LTR-RTs were retained. In addition, false positives were discarded by the LTR_retriver pipeline[7]. After removing non-LTR repeat elements and tandem repeats, the full-length LTR retrotransposons with optimal frames were translated into amino acids. Their functional domains were predicted using HMMER[8] based on the Pfam database. Paralogs of the RT domain, specific for the Copia and Gypsy superfamilies, were then detected by the functional domain orders.

The RT protein sequences were aligned using MUSCLE (v3.8.31)[9], and maximum likelihood trees were built using FastTree[10] for the Copia and Gypsy superfamilies. The two ends of these LTR retrotransposons were aligned using MUSCLE[5,6], and the nucleotide distance (D) was estimated using the Kimura two-parameter (K2p) criterion[11], which was implemented in the distmat program of the EMBOSS package (v6.6.0)[12]. The rates of nucleotide substitution ($\mu$) were obtained from a previous study[13]. The insertion times of an LTR retrotransposon were calculated using $T = D/2\mu$.

**Gene expression level analysis**

RNA-seq reads were aligned to the Taxus genome using HISAT2 (v2.1.0)[14] with default parameters. The sequence alignment files generated by HISAT2 were subsequently inputted to StringTie (v1.3.5)[15] to generate counts of uniquely mapped reads of annotated genes from the reference annotation file. The genes that differed significantly between two tissues were identified with count-based methods in R packages edgeR (v3.18.1)[16]. Benjamini-Hochberg correction was used to correct for multiple comparisons (with a false discovery cut-off $< 0.05$). The read counts of genes were transformed to FPKM values using fragments mapping per kilobase of exon per million fragments mapped. The FPKM values were used to represents the expression level of genes. Whereas, the genes with high sequence similarity were difficult to separate the reads mapping on these genes based on the methods above. To detected the high similarity sequence genes, we adjusted the threshold (the min acceptable alignment score parameter --score-min set as L,0, 0 in HISAT2) to no mismatch and get the alignment files. The expression level of genes was calculated using StringTie (v1.3.5)[15] using the alignment file as input files. Then, the relative low nucleotide differences distributed on the exons could use to distinguish the different expression level

**Ortholog analysis**

Publicly available genome and annotated peptide sequences of *A. thaliana* (TAIR 10), *O. sativa* (IRGSP-1.0), *A. trichopoda* ( AMTR1.0), and *S. moellendorffii* (v1.0) were downloaded from EnsemblPlants (http://plants.ensembl.org/), while the *P. abies* (v1.0b) and *G. biloba* (v1.0) sequences were downloaded from TreeGenes[17]. An all-to-all blast was performed using blastall[18] and the data were then processed to find orthologous gene groups using OrthoMCL[19]. Subsequently, 193 single-copy gene families (out of a total of 35,298 gene families) were found, and their protein sequences were aligned with MUSCLE[20]. The aligned sequences were then integrated into a single concatenated sequence for each species, and the conserved blocks were identified for further phylogenetic analysis using Gblocks[21]. The maximum likelihood phylogenetic tree was constructed using RAxML (Random Accelerated Maximum Likelihood)[22] with the GAMMAI+LG model, and the absolute rates of divergence times were estimated using r8s[23]. The number of expanded or contracted gene families along each branch was computed using CAFÉ[24] with a *p*-value cutoff of 0.05. Gene Ontology enrichment analysis was performed and visualized using clusterProfiler[25].

**Expression and purification of TSs in *E. coli***

To obtain recombinant TSs-His6, pET28b::*TS1* and pET28b::*TS2* were transformed into *E. coli* BL21 (DE3) and streaked on LB plates with kanamycin (50 μg $mL^{-1}$). A single positive colony was cultured at 37°C overnight in 5 mL of LB liquid medium with the same kanamycin concentration. The overnight cultures were used for expanding the culture in 500 mL of fresh medium and grown until the $OD_{600}$ reached 0.4, at which point, 1 mM isopropyl $\beta$-D-1-thiogalactopyranoside (IPTG) was added to induce expression. Expression induction was performed in a 180 rpm shaker at 16°C overnight, and harvested cells were resuspended in lysis buffer (50 mM sodium phosphate, pH 8.0 and 300 mM NaCl and 10 mM imidazole). After cell lysis by sonication and centrifugation at 4,000 g for 20 min at 4°C, the total protein supernatant was loaded onto a 1.5 mL HisPur Ni-NTA resin-packed column (Thermo Scientific, U.S.A), and the recombinant TSs-His6 were finally eluted with elution buffer (50 mM sodium phosphate, pH 8.0, 300 mM NaCl and 250 mM imidazole). The purified recombinant TSs-His6 were desalted into enzyme assay buffer (25 mM HEPES, pH 8.5, 10% glycerol, 5 mM DTT, 5 mM sodium ascorbate, 5 mM sodium metabisulfite, and 1 mM $MgCl_2$) and concentrated by centrifugation with a 30-kDa cutoff Millipore concentrator. The content and purity of the recombinant TSs-His6 were determined by a BCA protein assay kit (Beyotime, China).

#### *In vitro* characterization of T5αH1 and its homologous genes

The ORFs of Taxus *T5αH1* (*ctg5306_gene.3*), *T5αH2* (*ctg7747_gene.2*), *T5αH3* (*ctg2768_gene.2*), 55326109 (*ctg7747_gene.3*), and 55305455 (*ctg7747_gene.4*) were cloned from the Taxus cell line cDNA and inserted into the yeast expression vector pESC-His. Thus, the recombination plasmid pESC-His-*CYP725As-Flag* was obtained after positive clone screening. pESC-His-*CYP725As-Flag* was expressed in the yeast WAT11 strain[26]. Their expression was confirmed by western blot assay. All expression plasmids were constructed using the Hieff Clone™ One Step Cloning Kit (YEASEN, China) and primers used in this work are given in Supplementary Table 29.

For the *in vitro* enzyme assay, microsomes of the yeast strain WAT11 expressing pESC-His-CYP725As-Flag were prepared as described previously[27]. As a control, microsomes of the WAT11 strain harboring the empty vector pESC-His were prepared. The activity of the pESC-His-CYP725As-Flag protein was tested in a 1 mL mixture containing 50 mM HEPES buffer (pH 7.5), 100 µM taxadiene substrate, 500 µM NADPH, and 2 mg of pESC-His-CYP725As-Flag-containing microsomal proteins and overlaid with 500 µL of pentane. As a control, microsomes harboring the empty vector pESC-His were prepared. After incubating the mixture at 32°C for 2 h, the pentane layer was separated for GC/MS analysis.

The previous peroxide and stability assays indicated that OCT could be oxidized from taxa-4(5),11(12)-diene by hydrogen peroxide, and OCT can spontaneously form iso-OCT at 37°C overnight[28]. For given OCT and iso-OCT, 100 µL of the above purified taxadienes (approximately 200 ug $L^{-1}$) were concentrated by $N_2$ gas in a glass GC vial, then the residue was resuspended in 500 µL of hydrogen peroxide and shaken at 37°C and 220 rpm overnight. After reacting overnight, the taxanes were extracted with hexane and analyzed with GC-MS.

#### Transcription analysis under methyl jasmonate (MeJA) treatment

The *T. chinensis* var. *mairei* LC cell lines were divided into two groups: the test group was treated with 100 µM MeJA solution, whereas the control group was treated with solvent (ethanol). Samples were harvested after 0, 2, 4, 8, and 24 h of elicitation for gene expression analysis and taxane measurements. Experiments were carried out in triplicate. Total RNA was isolated from the MeJA-treated samples using the EASYspin Plant RNA isolation kit (Aidlab, Beijing, China), and 1 µg of cDNA was synthesized using Hifair® III 1st Strand cDNA Synthesis SuperMix for qPCR (gDNA digester plus) (YEASEN, China). Transcript abundance was measured using a QuantStudio™ 3 System with Hieff® qPCR SYBR® Green Master Mix (Low Rox Plus) (YEASEN, China). All real-time PCRs were repeated with at least two technical and three biological replicates. Mean cycle threshold (*Ct*) values were normalized using *T. chinensis* var. *mairei* actin 1 gene (*7G702435613*) as a validated reference gene. The relative gene expression value was calculated using the $2^{-\Delta\Delta Ct}$ method. Gene-specific primers are listed in Supplementary Table 29.

**Metabolite extraction and chromatography-mass spectrometry analysis**

The cell samples were extracted with a modified Wolfender method. One hundred milligrams of freeze-dried cell powder was weighed and transferred into a 2-mL centrifuge tube with 10 µL internal standards (500 ng $mL^{-1}$ dexamethasone). Then, the samples were resuspended in 1.5 mL extraction buffer (methanol:water = 80:20, v/v), and a 30 min ultrasonic-assisted extraction process was applied. After 15 min of centrifugation at 14,000 g, the supernatants were dried in a LABCONCO CentriVap vacuum centrifugal concentrator and resuspended in 200 µL methanol solvent (80:20, v/v).

GC/MS analysis was performed on an Agilent 7890B GC machine (Agilent Technologies, Waldbronn, USA) equipped with an Agilent 7000C mass selective detector at 70 eV and 1.2 mL $min^{-1}$ helium flow. One to five microliters of the sample was injected and analyzed on an Agilent HP-5MS column (5% phenyl methyl silox, 30 m × 250 µm internal diameter, 0.25-µm film thickness). The oven temperature program was as follows: 45°C for 1 min, then a 10°C $min^{-1}$ ramp to 250°C, and a hold at 250°C for 5 min. The injection temperature was 250°C. Full mass spectra were generated for metabolite identification by scanning the mass-to-charge ratio (*m/z*) range from 40 to 350.

Liquid chromatography-mass spectrometry (LC-MS) analysis was carried out on an ACQUITY UPLC I-Class (Waters) coupled to a 4500 QTRAP triple quadrupole mass spectrometer (AB SCIEX) equipped with a 50 × 2.1 mm, 1.8 μm ACQUITY UPLC™ Waters Xselect HSS T3 column (Waters). Ten microliters of supernatant was loaded each time and then eluted at a flow rate of 200 μL min$^{-1}$ with initial conditions of 40% mobile phase A (0.1% formic acid in methanol) and 60% mobile phase B (0.1% formic acid in water) followed by a 10-min linear gradient to 100% mobile phase B. The autosampler was set at 10°C. Mass spectrometry was operated separately in positive electrospray ionization mode. The analyte [M+H] was selected as the precursor ion. The quantitation mode was multiple reaction monitoring (MRM) mode using mass transitions (precursor ions/product ions). The temperature of the ESI ion source was set at 500°C. The curtain gas flow was set at 20 psi, collisional activated dissociation (CAD) gas was set as the medium, and the ion spray voltage was (+) 5,500 V, with ion gases 1 and 2 set as 50 psi. AB SCIEX Analyst 1.6.3 Software (Applied Biosystems) was used for data acquisition and processing.

**2. Supplementary Figures 1-11**

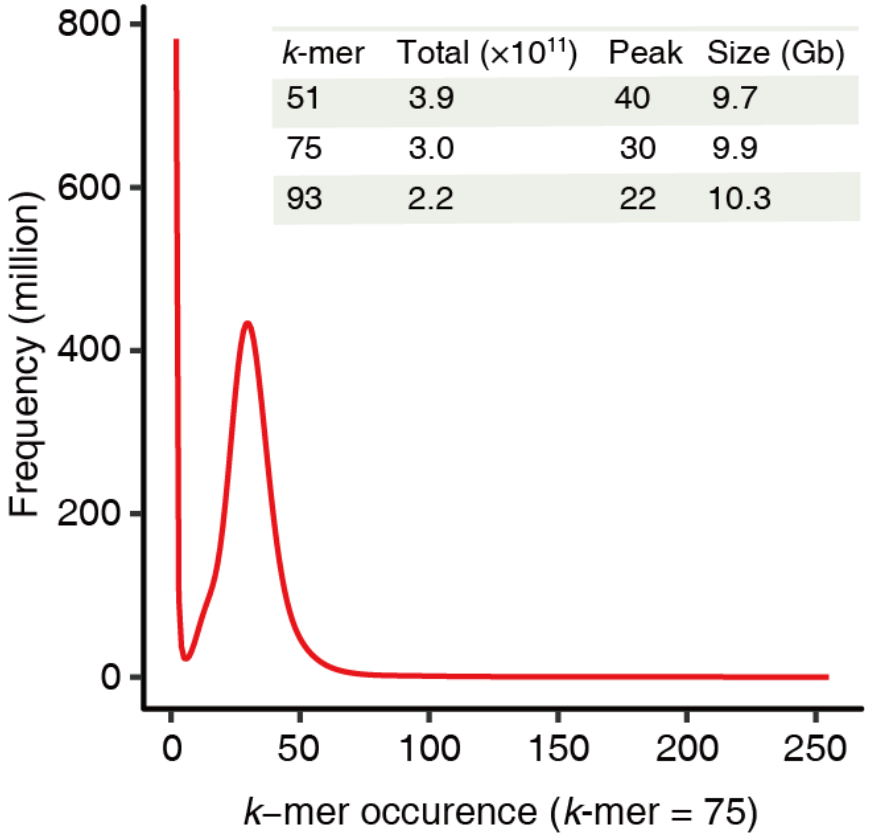


**Supplementary Fig. 1 Genome size estimation of *T. chinensis* var. *mairei* based on *k*-mer distribution.** The X-axis represents the occurrence of *k*-mers, and the Y-axis represents the frequency. The *k*-mer values for different genome sizes are shown in the inner table.

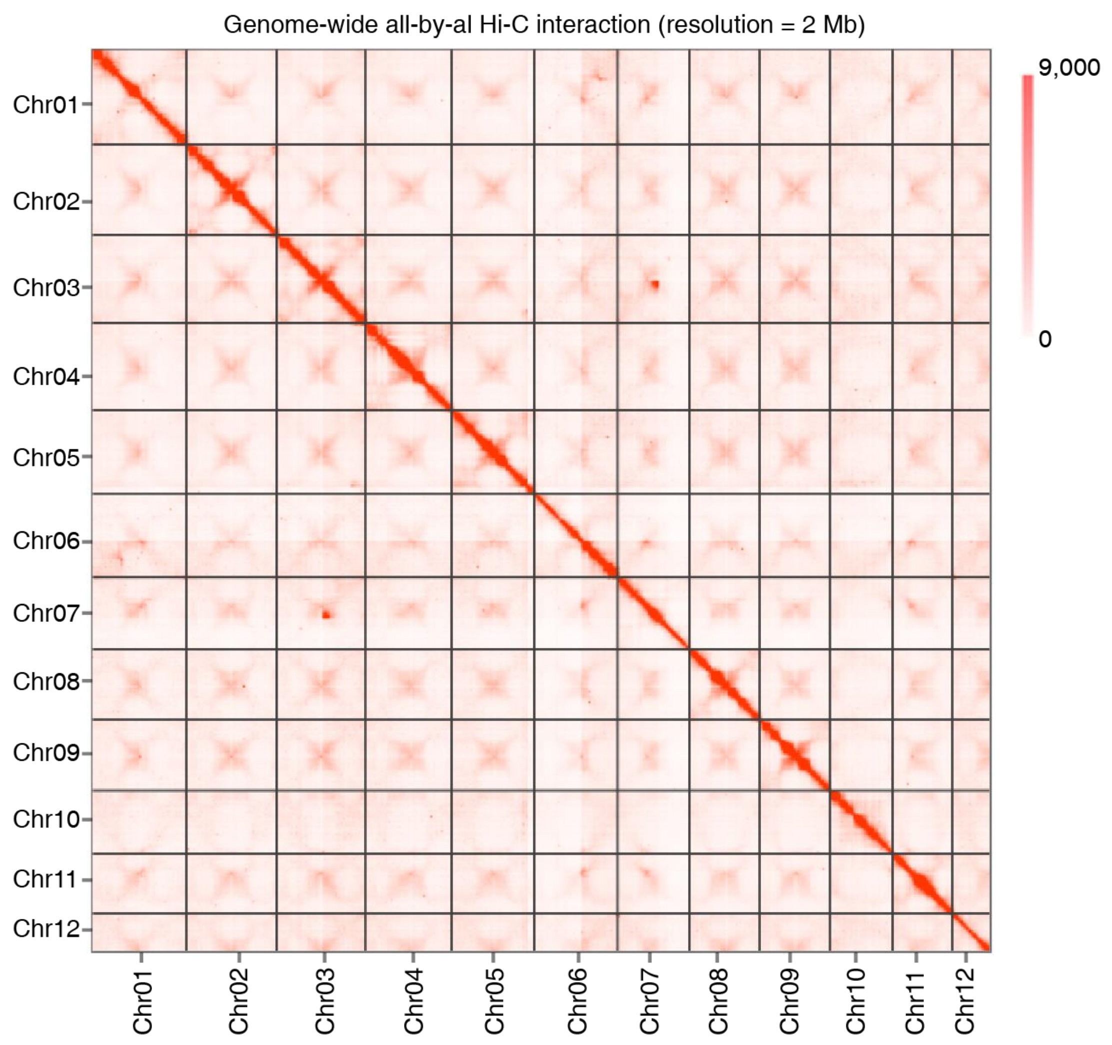


**Supplementary Fig. 2 Genome-wide all-by-all Hi-C interaction.** The heat map shows Hi-C interactions under a resolution of 2 Mb. Darker red pixels indicate higher contact probabilities. The number on the scale bar indicates the number of links after logarithmic analysis.

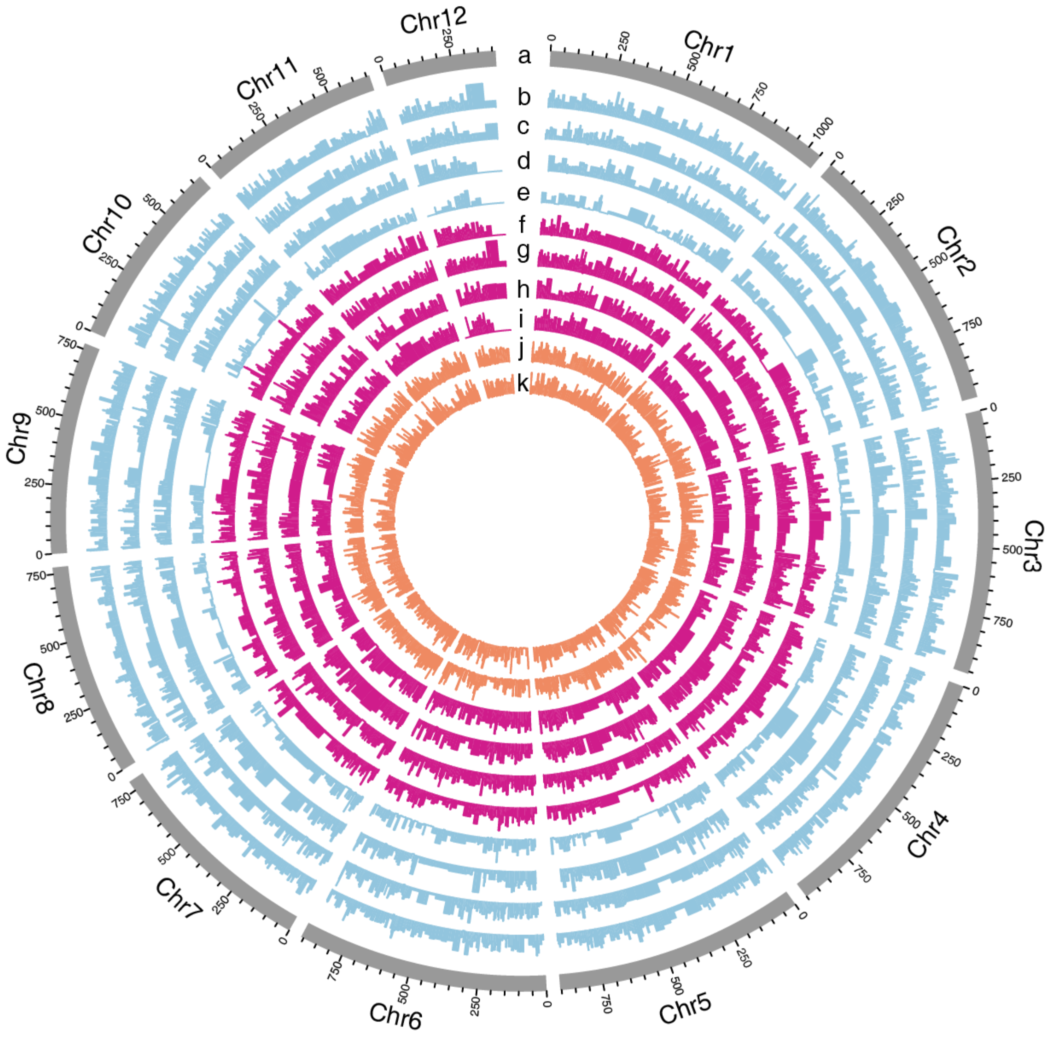


**Supplementary Fig. 3 Genomic landscape of the twelve pseudochromosomes.** Track a represents the length of the pseudochromosomes (Mb); b, c, d, and e show the expression of tissue-specific genes in the bark of stem, root, strobili and leaf from the male Taxus plant, respectively; f, g, h, and i show the expression of tissue-specific genes in the bark of stem, root, strobilus and leaf from the female Taxus plant, respectively; j and k display high- and low- producing paclitaxel cell lines, respectively.

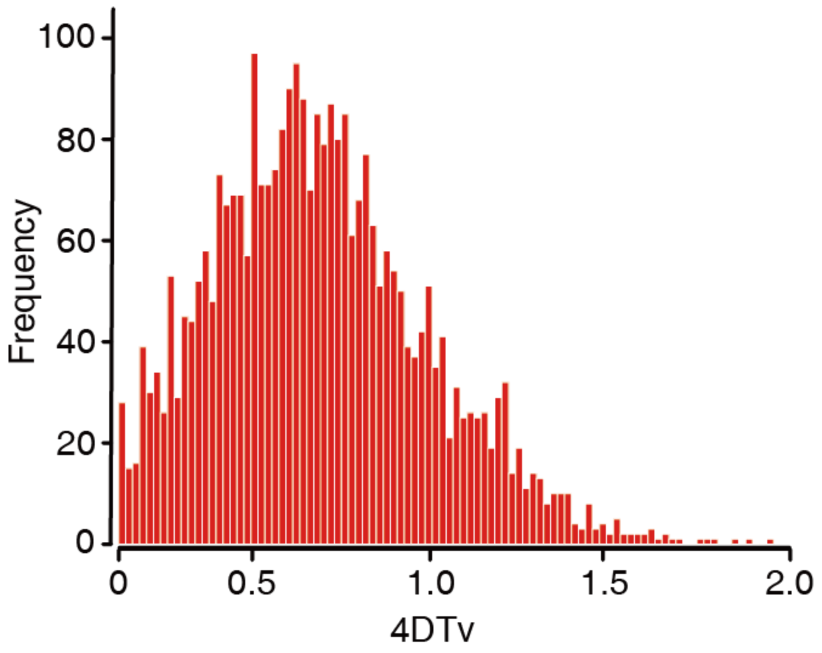


**Supplementary Fig. 4 Whole genome duplication (WGD) analysis based on the substitution rate distribution of paralogs.** The 4DTv values of paralogs were calculated using *KaKs*_calculator with the YN model. The X-axis is the value of fourfold synonymous third-codon transversions (4DTv) for paralogous pairs in the Taxus genome, and the Y-axis represents the frequency.

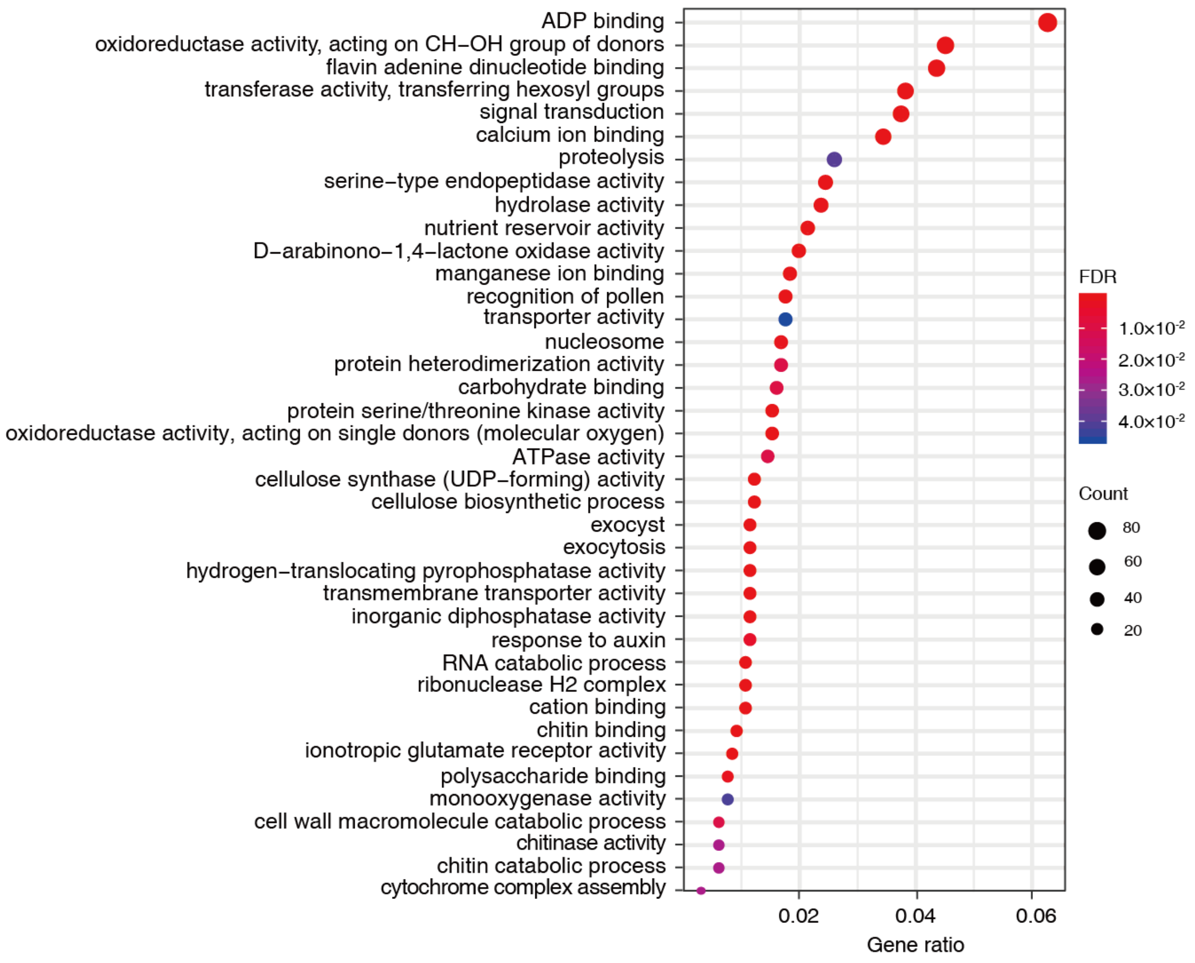


**Supplementary Fig. 5 Gene Ontology (GO) enrichment for gene families with significant expansion.** GO enrichment analysis of a subset of 142 gene families with significant expansion (*p* < 0.05); FDRs were adjusted for multiple testing. The size and color of dots indicate the number of genes and false discovery rate (FDR), respectively. The X-axis represents the gene ratio, and the GO terms are listed on the Y axis.

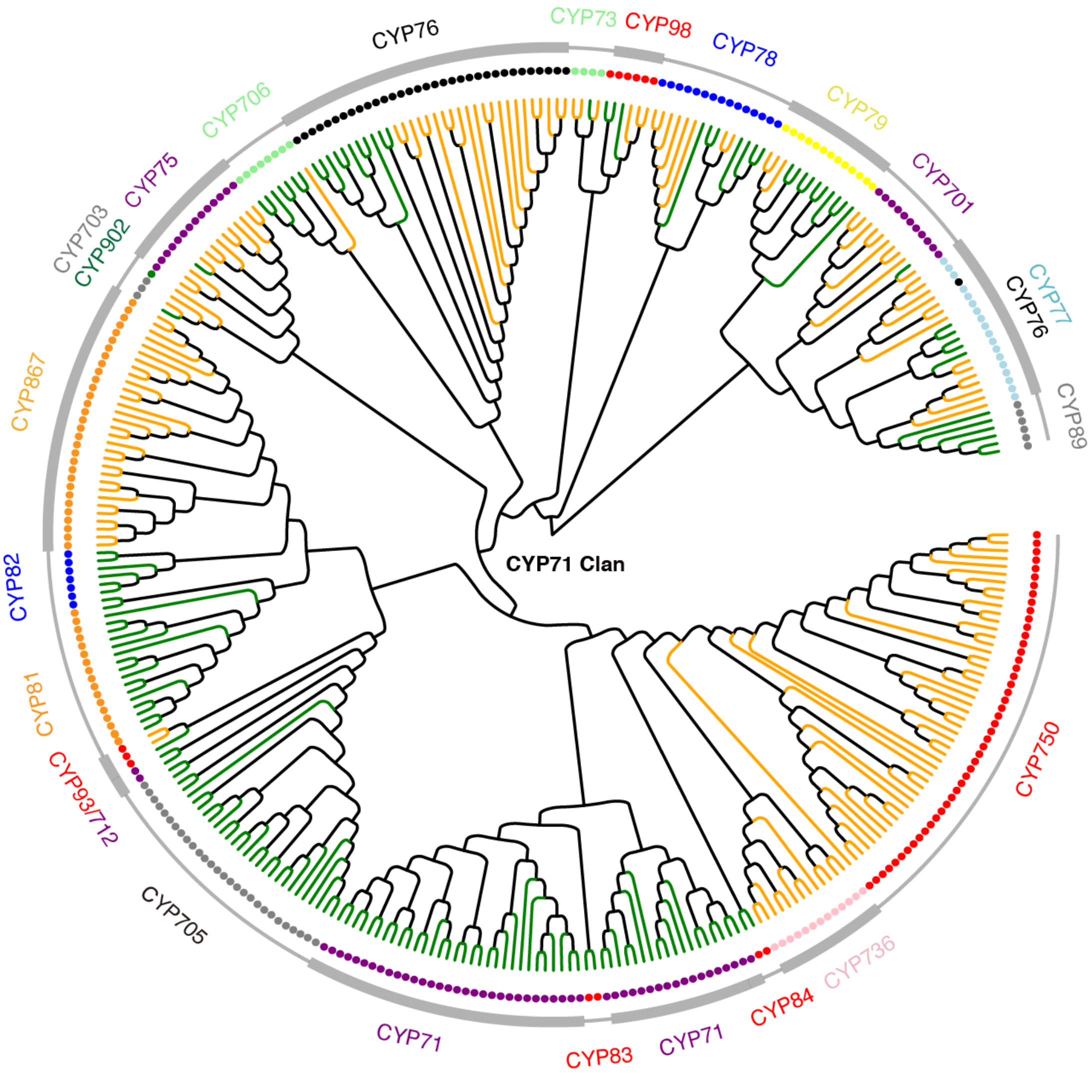


**Supplementary Fig. 6 Phylogenic analysis of A-type CYP450 families.** The green and orange branches indicate the sequences from Arabidopsis and *T. chinensis* var. *mairei*, respectively. The dots represent CYP450 genes. The outermost circle indicates the CYP450 gene family.

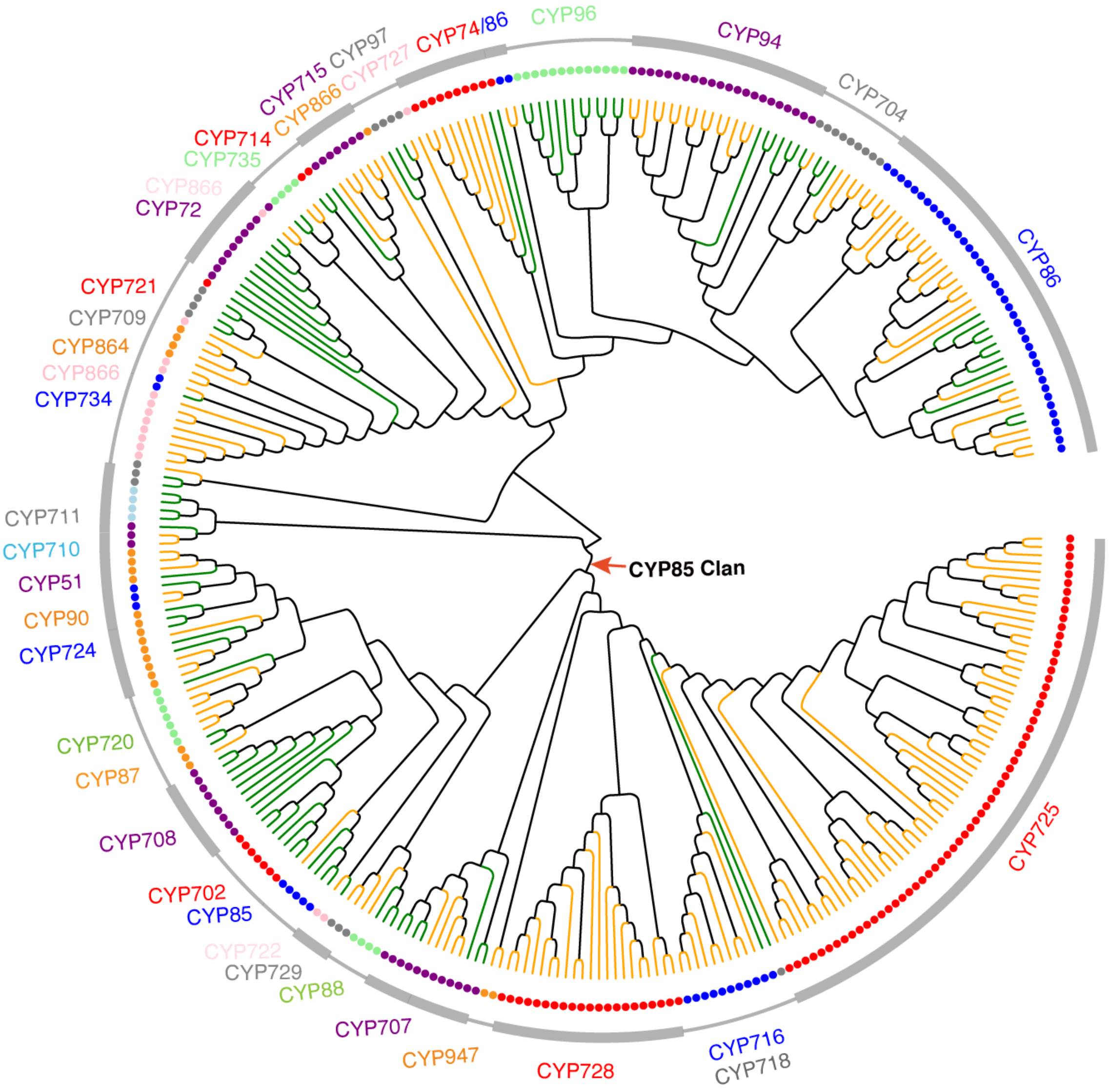


**Supplementary Fig. 7 Phylogenic analysis of non-A-type CYP450 families.** The green and orange branches indicate the sequences from Arabidopsis and *T. chinensis* var. *mairei*, respectively. The dots represent CYP450 genes. The outermost circle indicates the CYP450 gene family.

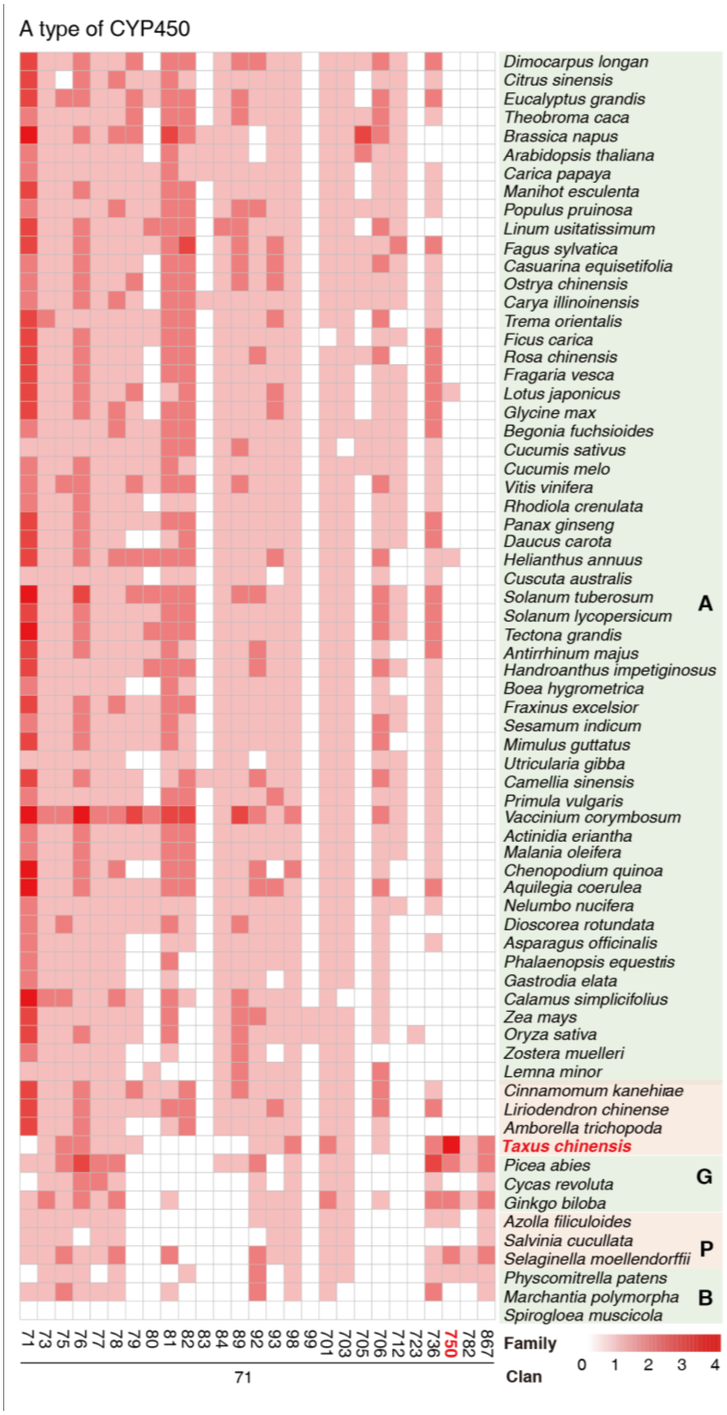


**Supplementary Fig. 8 Heat map of the number of A-type CYP450 genes in 69 representative plant species.** Each gene family is represented as a square, with the red color representing the number of genes in the corresponding family. The depth of the red color is divided into five levels, namely, 0, 1, 2, 3, and 4, which correspond to 0, 1-10, 10-50, 50-100 and more than 100 genes, respectively. A, Angiosperms; G, Gymnosperms; P, Pteridophytes; and B, Bryophytes.

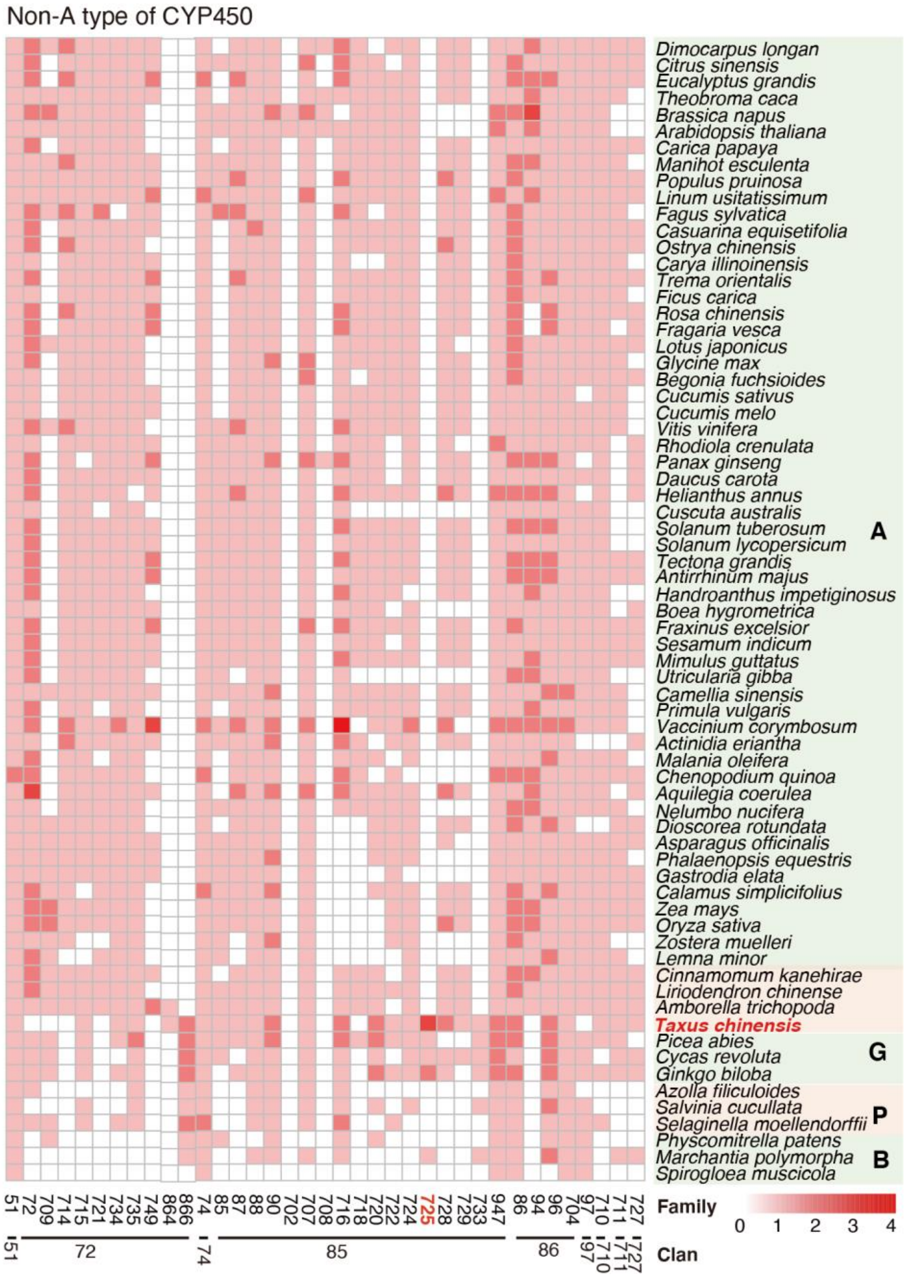


**Supplementary Fig. 9 Heat map of the number of non-A-type CYP450 genes in 69 representative plant species.** Each gene family is represented as a square, with the red color representing the number of genes in the corresponding family. The depth of the red color is divided into five levels, namely, 0, 1, 2, 3, and 4, which correspond to 0, 1-10, 10-50, 50-100 and more than 100 genes, respectively. The family or clan name of non-A-type CYP450 genes is marked below the heat map. A, Angiosperms; G, Gymnosperms; P, Pteridophytes; and B, Bryophytes.

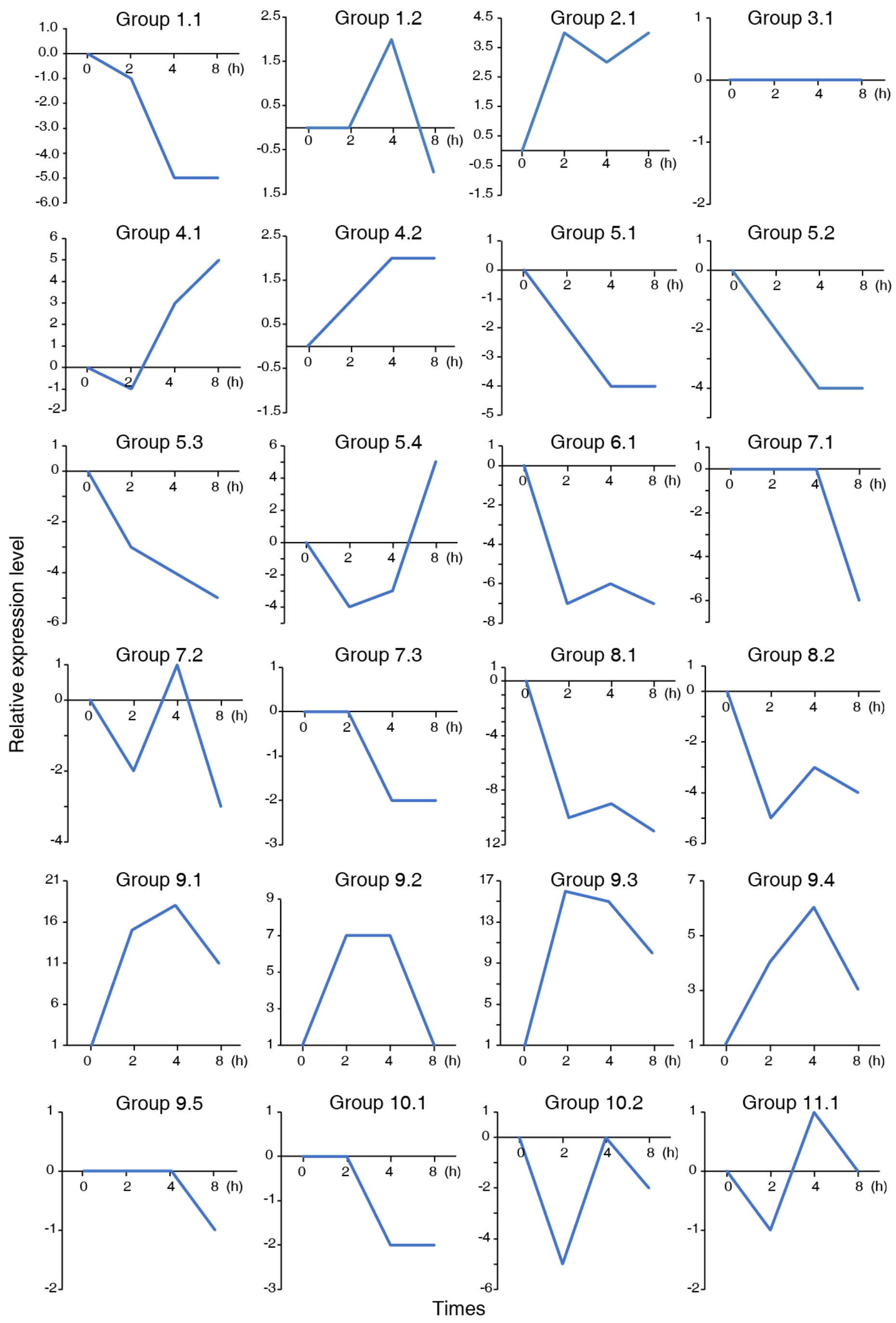


**Supplementary Fig. 10 Group-based gene expression profiles in response to MeJA treatment.** RNA sequencing analysis was performed with the low paclitaxel-yielding cell line (LC) treated with 100 μM MeJA or 0.5% EtOH solution for 0, 2, 4, and 8 h. The expression of the gene group was calculated by summing the expression levels of each CYP450. Each upregulated and downregulated CYP450 was calculated as 1 and -1, respectively, based on their FPKM (Fragments Per Kilobase of transcript per Million mapped reads) values.

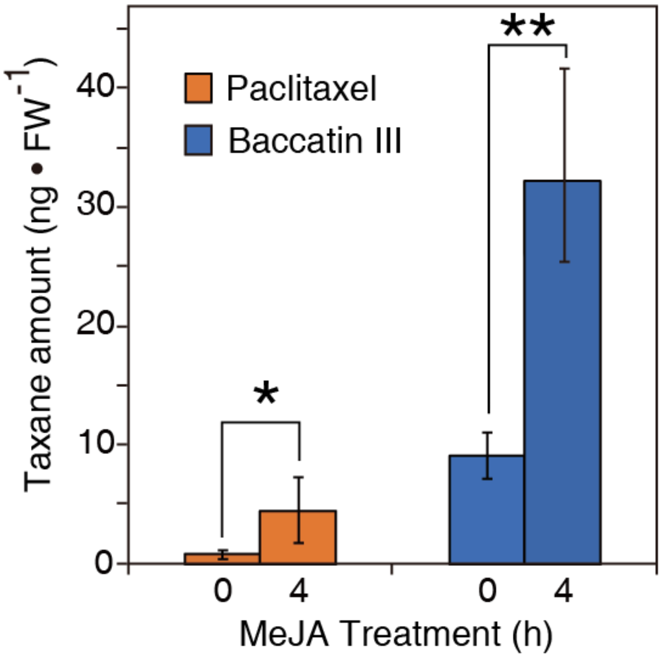


**Supplementary Fig. 11 The accumulation of paclitaxel and baccatin III in taxus cell lines induced by 100 μM MeJA at 0 hours and 4 hours.** The amounts of paclitaxel and baccatin III were measured by LC-MS analysis. FW indicates fresh weight (100 mg); error bars display the standard error (n = 3 biological replicates); asterisks show significant differences with $*P \leq 0.05$ and $**P \leq 0.01$ by two-tailed Student's t test.

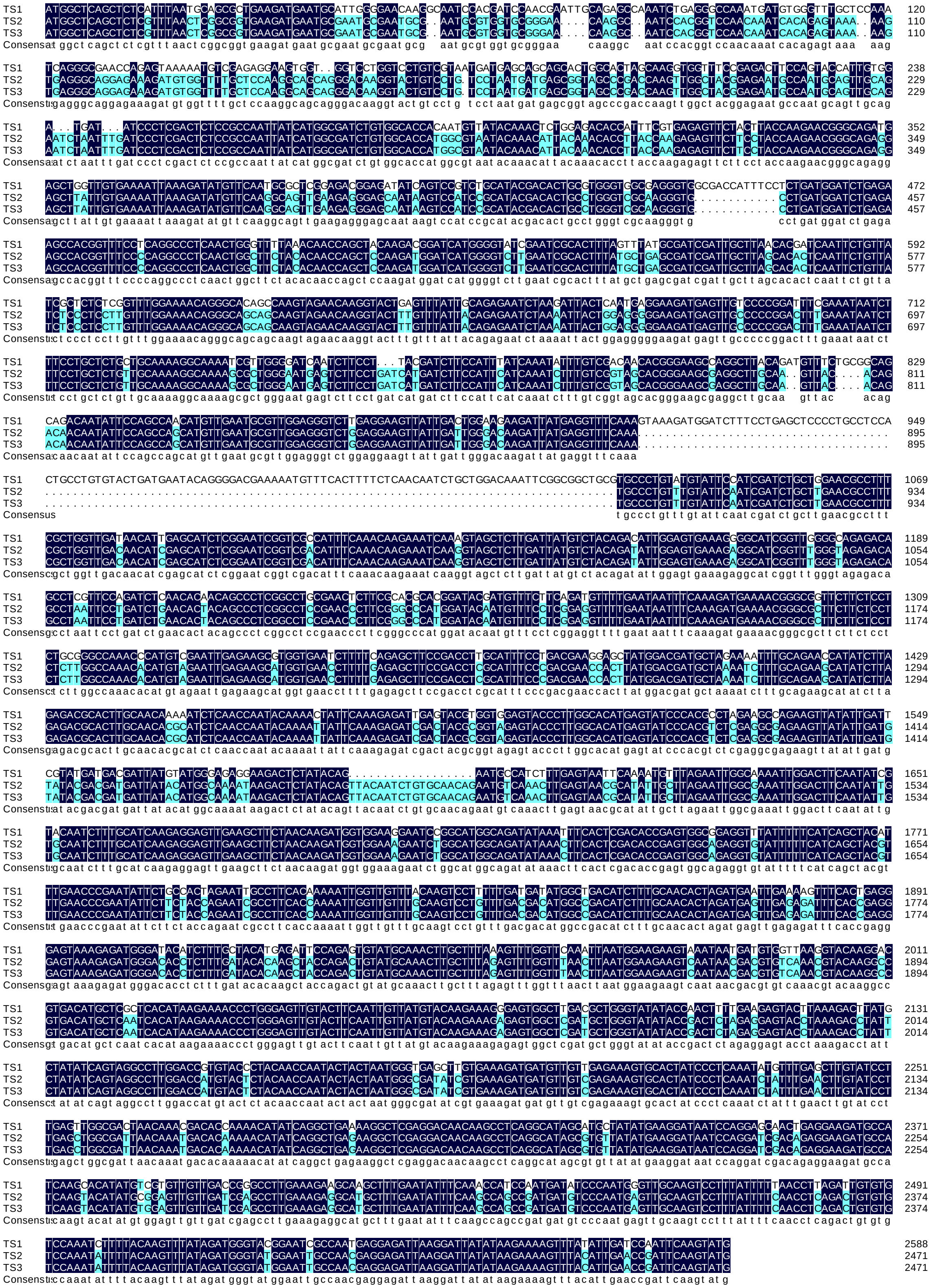

**Supplementary Fig. 12 Nucleotide sequence alignment of TSs.** TS1, 2, and 3 refer to *T. chinensis* var. *mairei* taxadiene synthase 1, 2, and 3, respectively.

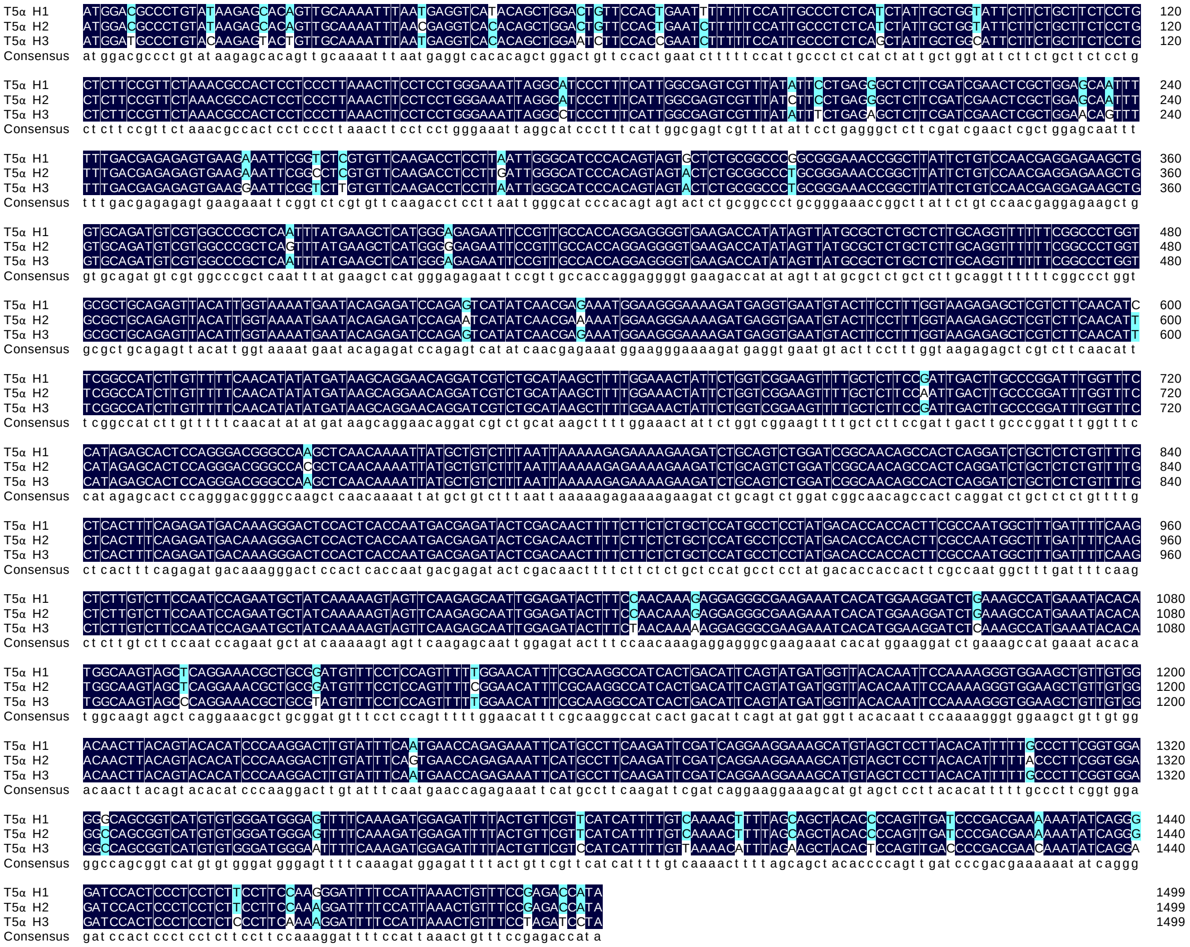

**Supplementary Fig. 13 Nucleotide sequence alignment of T5αHs.** T5αH1, 2, and 3 refer to *T. chinensis* var. *mairei* taxa-4(5),11(12)-diene-5α-hydroxylase 1, 2, and 3, respectively.

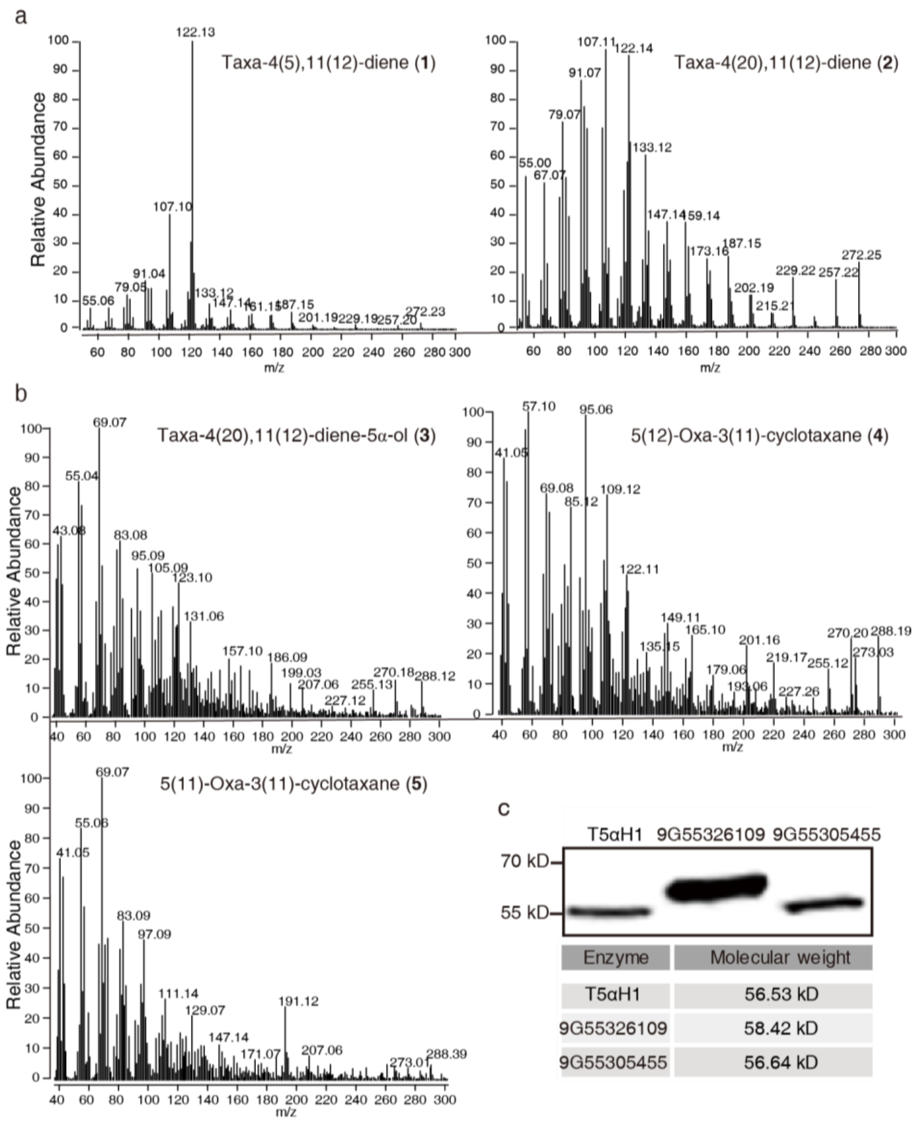


**Supplementary Fig. 14 Mass spectra of peaks (1, 2, 3, 4, and 5) shown in Fig. 3 and protein expression of the three CYP725As (T5αH1, 55326109, and 55305455) in yeast.** (**a**) Peaks 1 and 2 shown in Fig. 3b are the compounds taxa-4(5),11(12)-diene and taxa-4(20),11(12)-diene, respectively, produced by TSs; (**b**) peaks **3**, **4** and **5** shown in Fig. 3c represent the compounds taxa-4(20),11(12)-diene-5α-ol, OCT and iso-OCT, respectively, generated by T5αHs; (**c**) immunoblotting assays of three enzymes (T5αH1, 55326109 and 55305455) in the WAT11 strain. The mouse monoclonal antibody for the indicated epitope tags and sheep polyclonal antibody conjugated with horseradish peroxidase (Beyotime, China) were used as primary and secondary antibodies, respectively.

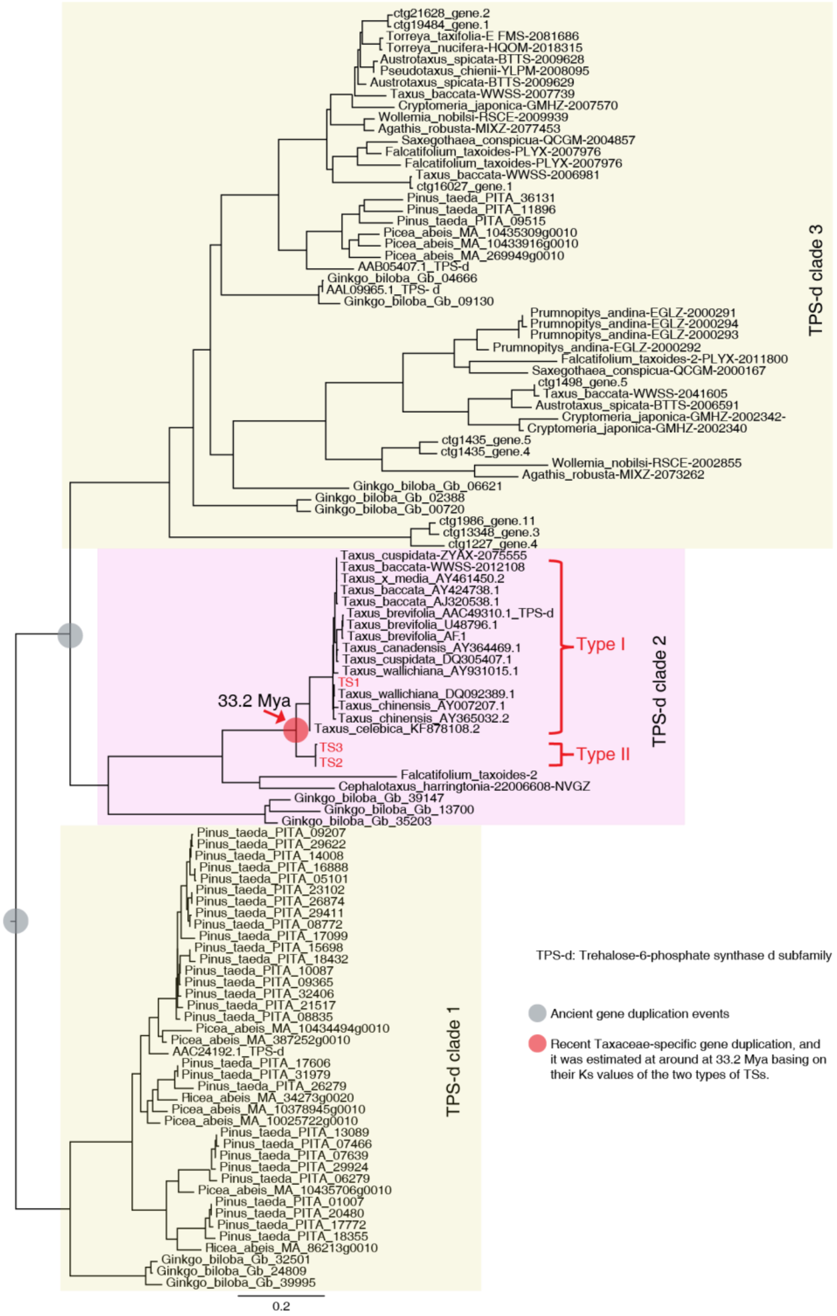


**Supplementary Fig. 15 Phylogenetic analysis of trehalose-6-phosphate synthase (TPS) genes from different plants.** The tree is generated from amino acid sequences by the maximum-likelihood method with 100 bootstraps. Ancient gene duplication events are indicated as gray dots, while the more recent Taxaceae-specific gene duplication is shown as a red dot. The *TS1/2/3* genes in *T. chinensis* var. *mairei* are highlighted in red.

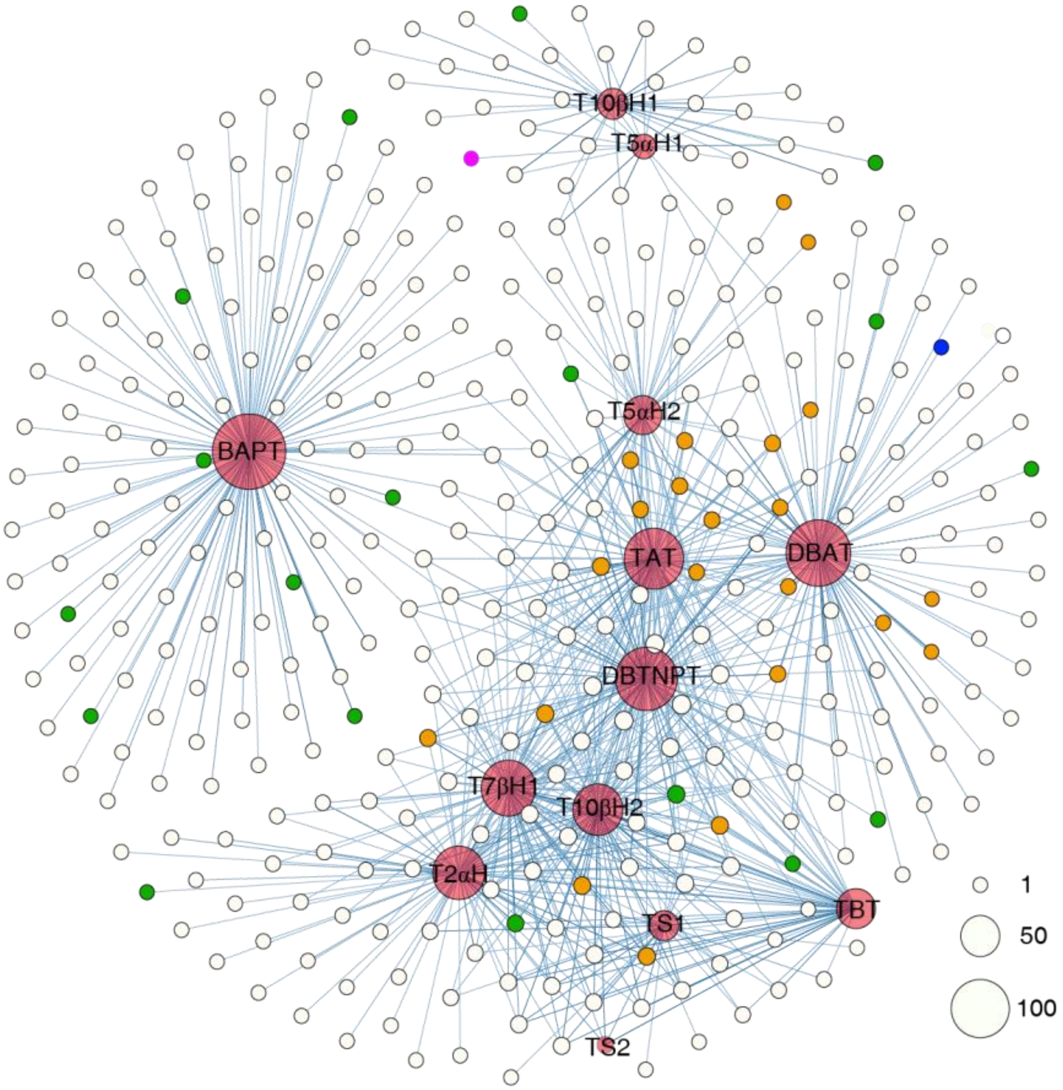


**Supplementary Fig. 16 Co-expression net of paclitaxel biosynthesis genes.** The genes with a Pearson correlation coefficient value above 0.75 are displayed on the net. The known paclitaxel biosynthesis genes, CYP725s, CYP450s, and the remaining genes are represented as red, orange, green, and white dots, respectively. The purple and blue dots show the two novel CYP725As 55305455 and 55326109, respectively. The size of the dot correlates with the gene number.

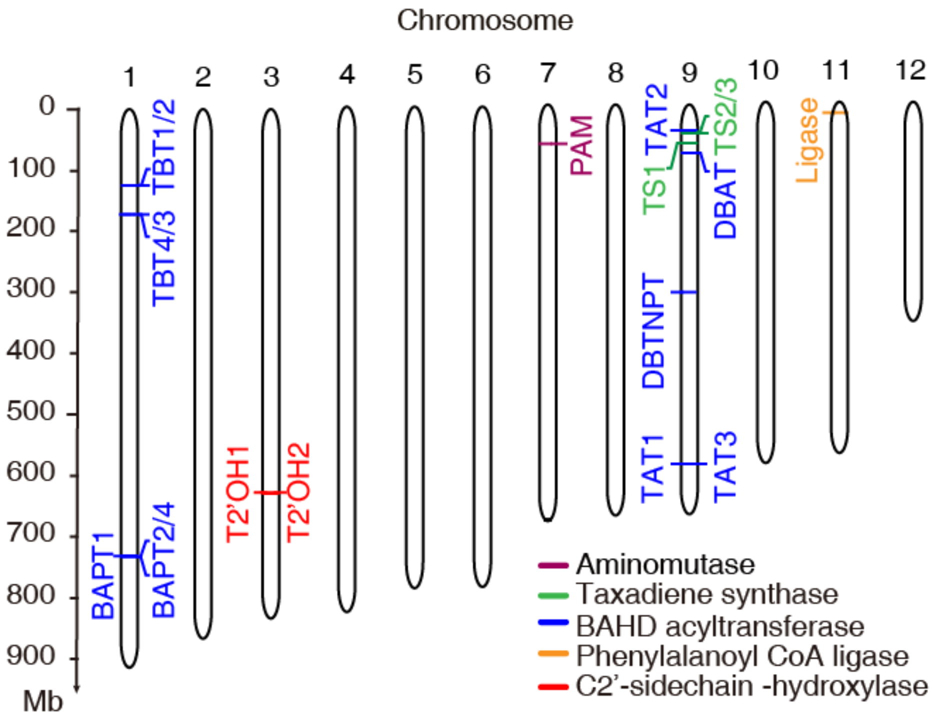


**Supplementary Fig. 17 Genomic location of the annotated genes known to be involved in paclitaxel biosynthesis, except for CYP450s.** The different colors of the short lines indicate the different types of annotated genes and their homologs in the paclitaxel pathway; the short purple, green, blue, orange, and red lines correspond to taxus aminomutase, taxadiene synthase, BAHD acyltransferase, ligase, and C2'-sidechain-hydroxylase.

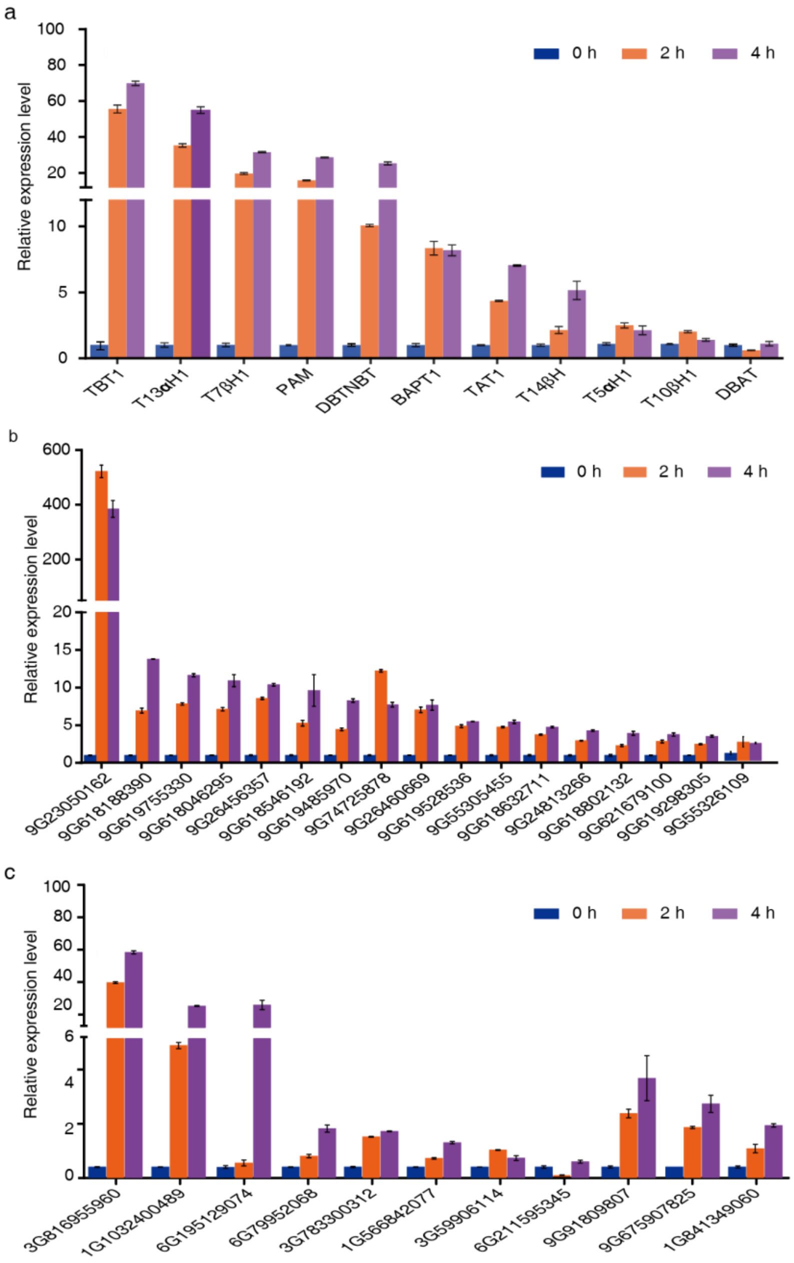


**Supplementary Fig. 18 Quantitative real-time PCR (qPCR) analysis of the gene expression of the candidate genes in the paclitaxel pathway.** The relative transcript abundance of the eleven defined paclitaxel biosynthetic genes (a), the sixteen CYP725A candidates (b), and the eight TFs and three BAHD acyltransferase genes (c) in MeJA-induced Taxus cell lines. The relative gene expression levels are represented as the average fold change ($2^{-\Delta\Delta Ct}$). The Taxus actin 1 gene (*7G702435613*) was used as an internal reference. Error bars indicate standard errors from three independent biological replicates.

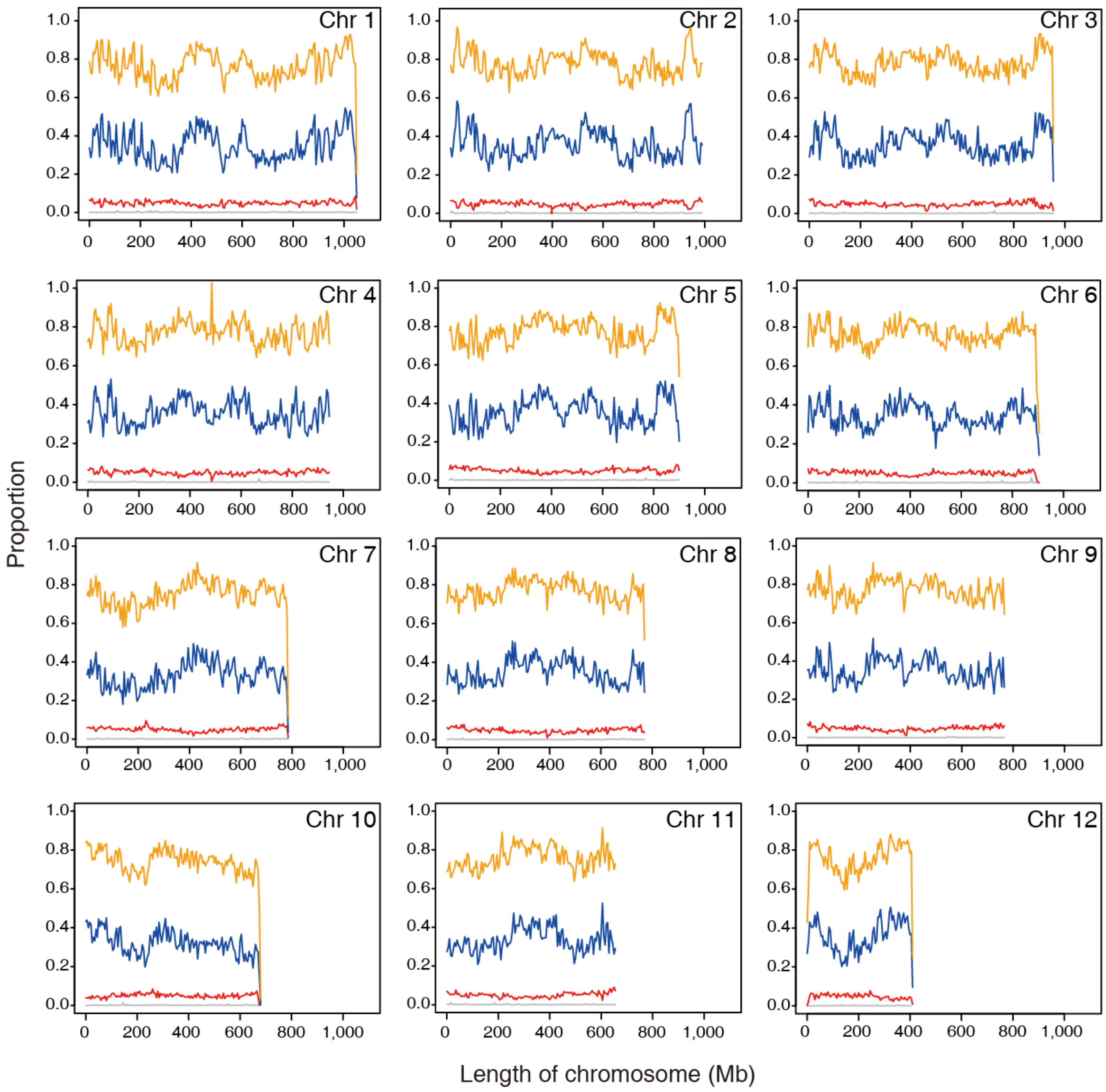


**Supplementary Fig. 19 Distribution of repeats and LTR on the chromosomes.** The lines indicate different elements (Orange: repeats; Blue: Gypsy; Red, Copia; Grey: Unknown LTR). Each point on the line represents the proportion of the component in the 5 Mb window.

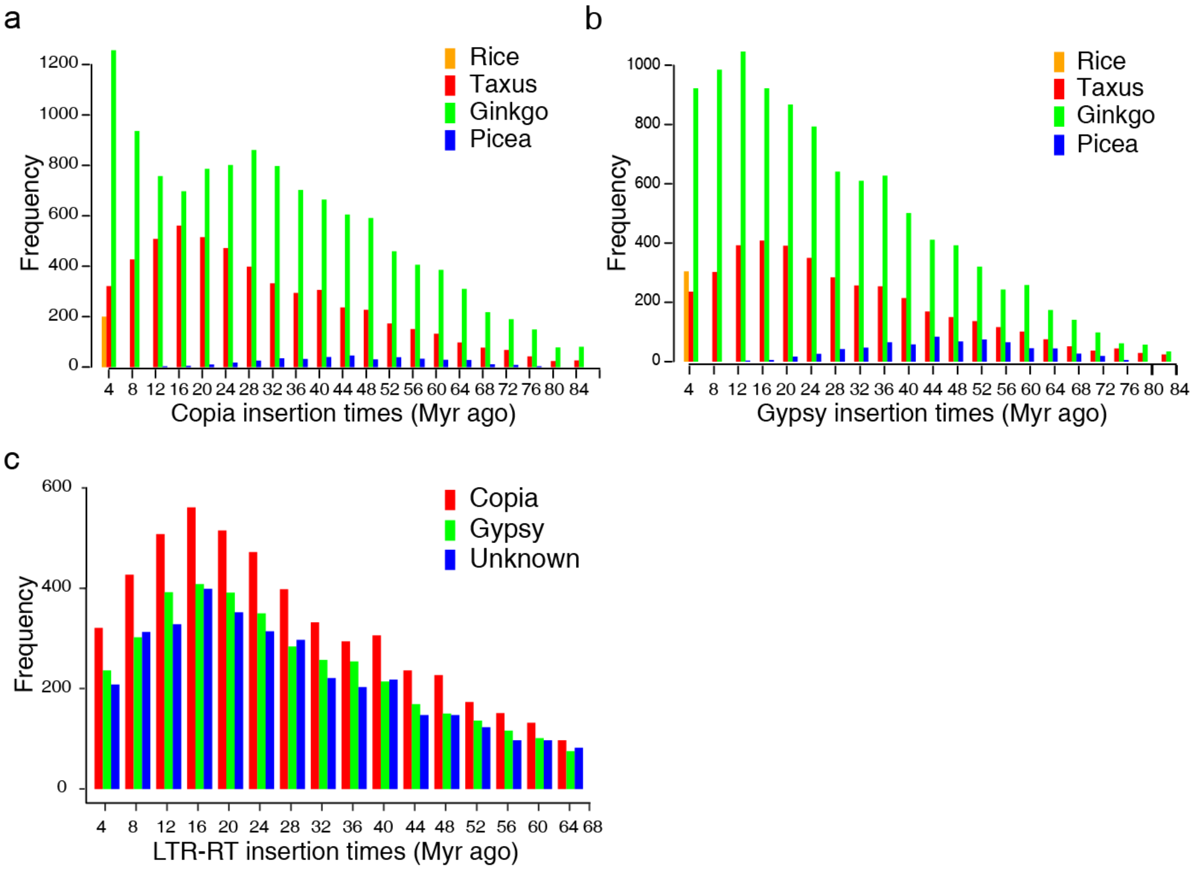


**Supplementary Fig. 20 Expansions and diverse sets of LTR elements in the Taxus genome.** a-b, The histogram shows the distributions of insertion times calculated for Gypsy and Copia in Taxus, ginkgo, picea, and rice. The different colors of the columns represent the Gypsy and Copia insertions in the four plants. c, The histogram shows the distributions of the insertion times calculated for the Taxus LTR elements (including Gypsy, Copia, and an unknown type).

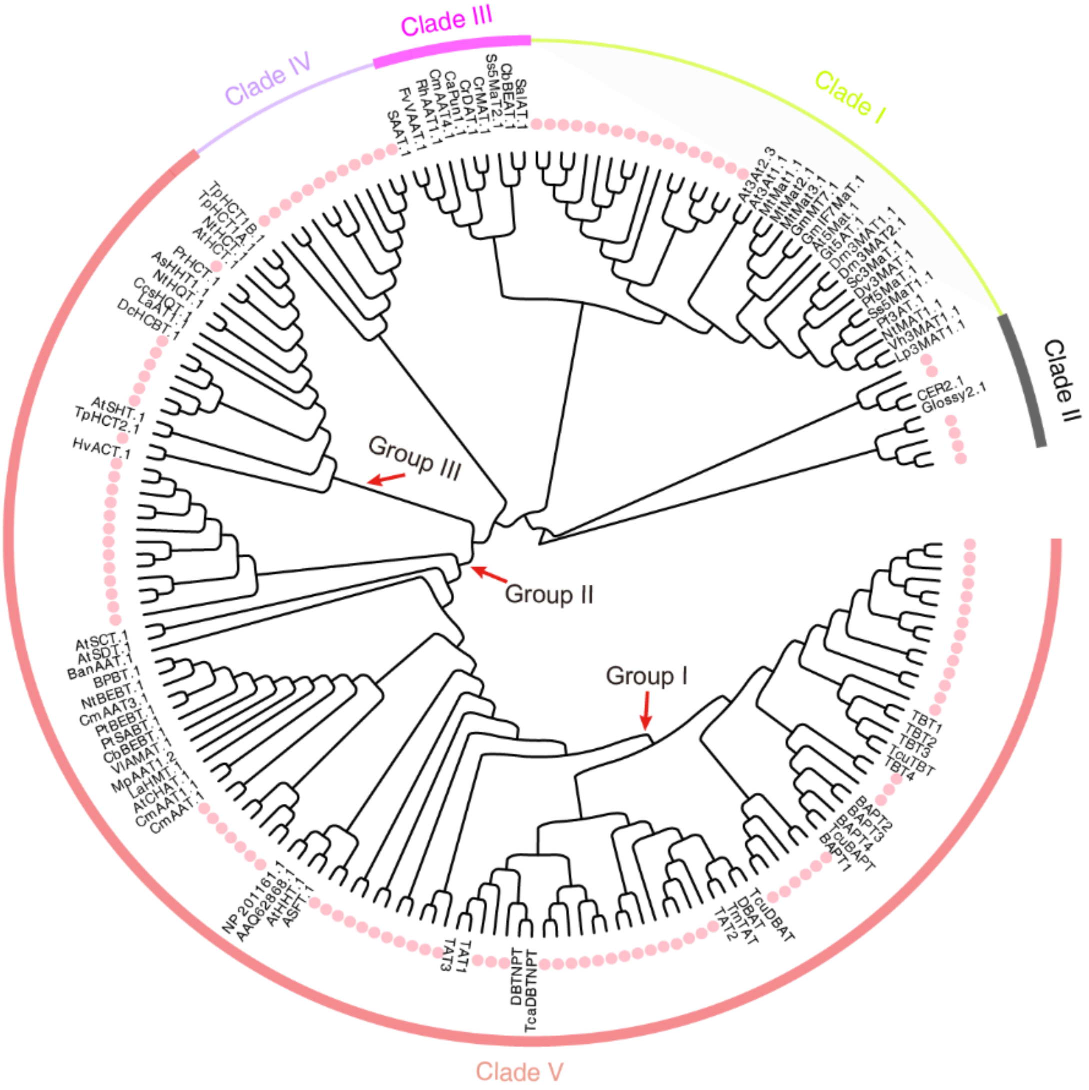


**Supplementary Fig. 21 Phylogenetic analysis of Taxus BAHD acyltransferase proteins.** The tree is generated using the neighbor-joining method and 1000 bootstraps based on putative amino acid full-length transferase sequences with MEGA 7.0 software and EvolView. The pink-red circle denotes transferase genes in *T. chinensis* var. *mairei*. The subgroups were designated based on a previous report[30]. All defined paclitaxel pathway BAHD acyltransferase genes are included in Group 1 of Clade V.

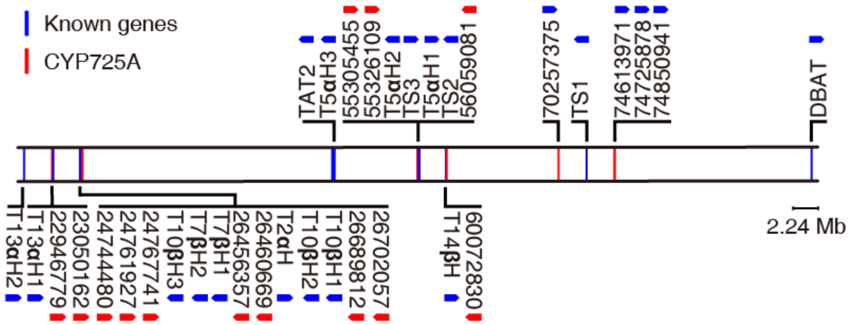


**Supplementary Fig. 22 Map of 17 defined genes and 18 novel CYP725As on chromosome 9.** The known genes in the paclitaxel biosynthesis pathway (known genes) are marked by blue lines, while the unknown CYP725As are shown in red lines. The arrows show gene orientations.

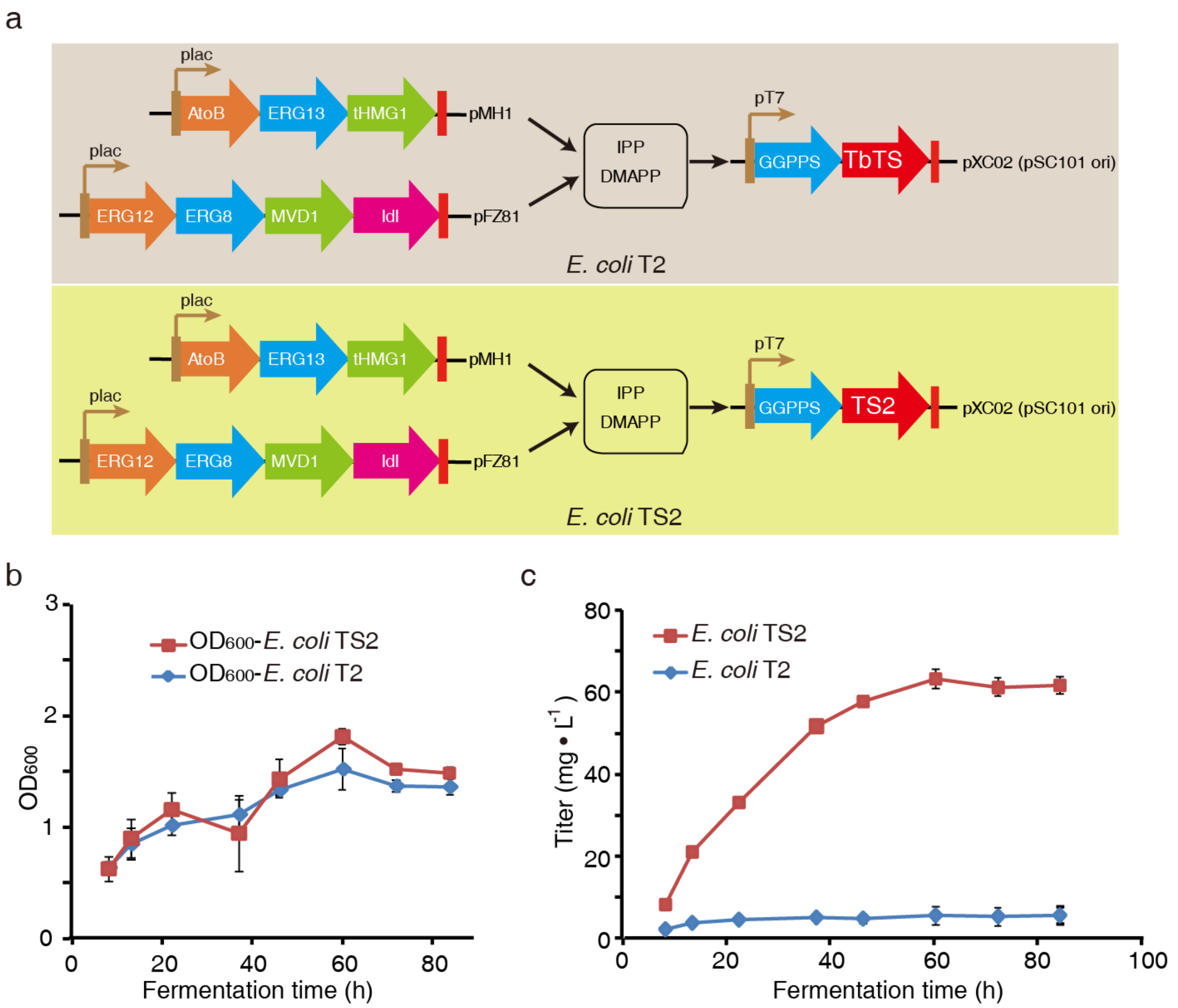


**Supplementary Fig. 23 Comparison of the two types of TS on the production of taxadiene in *E. coli*.** (a) Taxadiene-producing *E. coli* T2 (harboring pMH1, pFZ81, and pXC02) was constructed by coexpressing nine genes (*AtoB*, *ERG13*, *tHMG1*, *ERG12*, *ERG8*, *MVD1*, *IdI*, *GGPPS* and *TbTS*) in *E. coli*[29], while *E. coli* TS2 was generated by replacing *TbTS* with *TS2* based on the taxadiene-producing platform; (b) the cell concentrations of the strains *E. coli* T2 and *E. coli* TS2 were measured by $OD_{600}$ at set intervals (at 8, 13, 22, 37, 48, 60, and 72 hours); (c) the titers of taxadiene produced by *E. coli* T2 and *E. coli* TS2 in the shaking flask. TbTS, a *T. brevifolia* taxadiene synthase that shares 98.42 % amino acid sequence identity with TS1, represents type I TSs, while TS2, which shares 77 % protein sequence identity with TS1, is designated type II TSs. Error bars show standard error (n = 4 independent biological replicates).

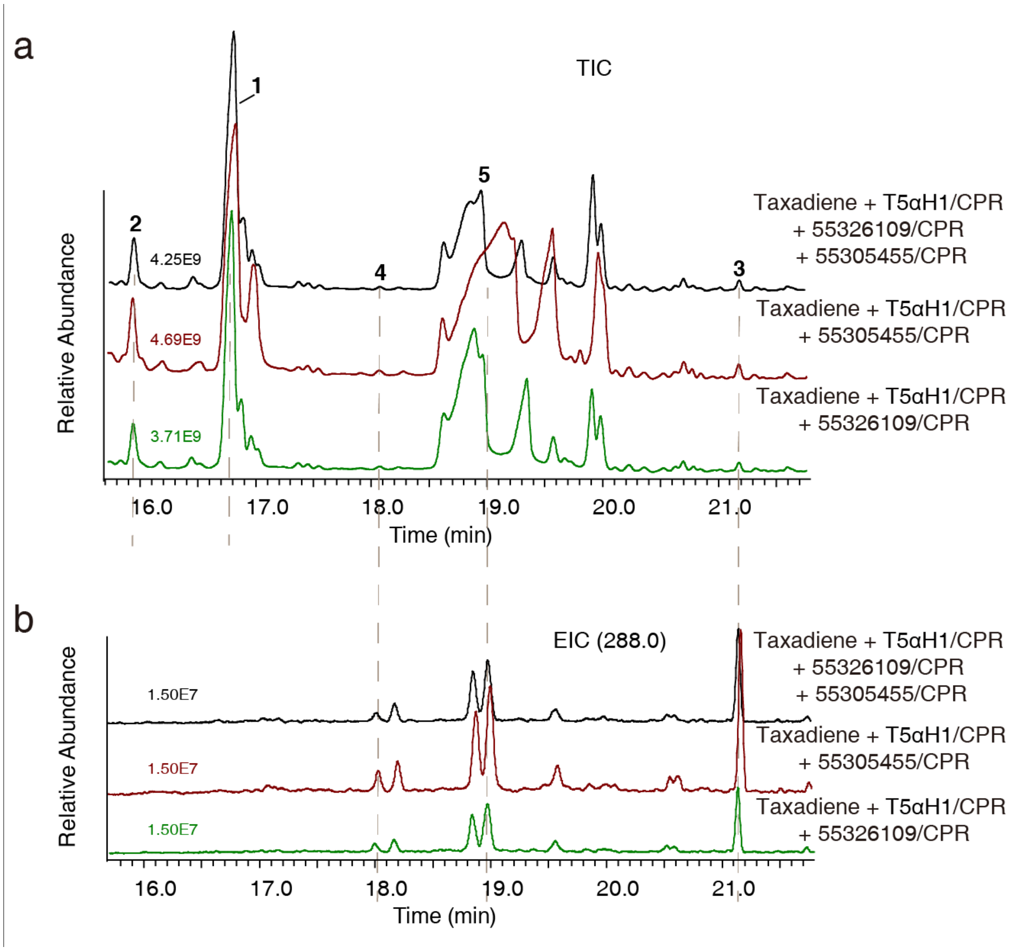


**Supplementary Fig. 24 Enzymatic analysis of CYP725As (55326109 and 55305455) with taxane as the substrate.** (a) Total ion chromatograms (TICs) of the *in vitro* assays. Peaks 1-5 are the substrates taxa-4(5),11(12)-diene, taxa-4(20),11(12)-diene, taxa-4(20),11(12)-dien-5α-ol, OCT and iso-OCT. The taxadiene substrates (peaks 1 and 2) were exogenously added, while the oxidized taxadienes (peaks **3**, **4** and **5)** were from the T5αH/CPR catalytic products. (b) Chromatograms of the extracted ions at 288 $m/z^+$ for the oxidized taxadienes (peaks 3, 4 and 5). There were no new peaks produced by the reaction mixtures, including T5αH/CPR and 55326109/CPR, T5αH/CPR and 55305455/CPR, and T5αH/CPR, and 55326109/CPR and 54359314/CPR.

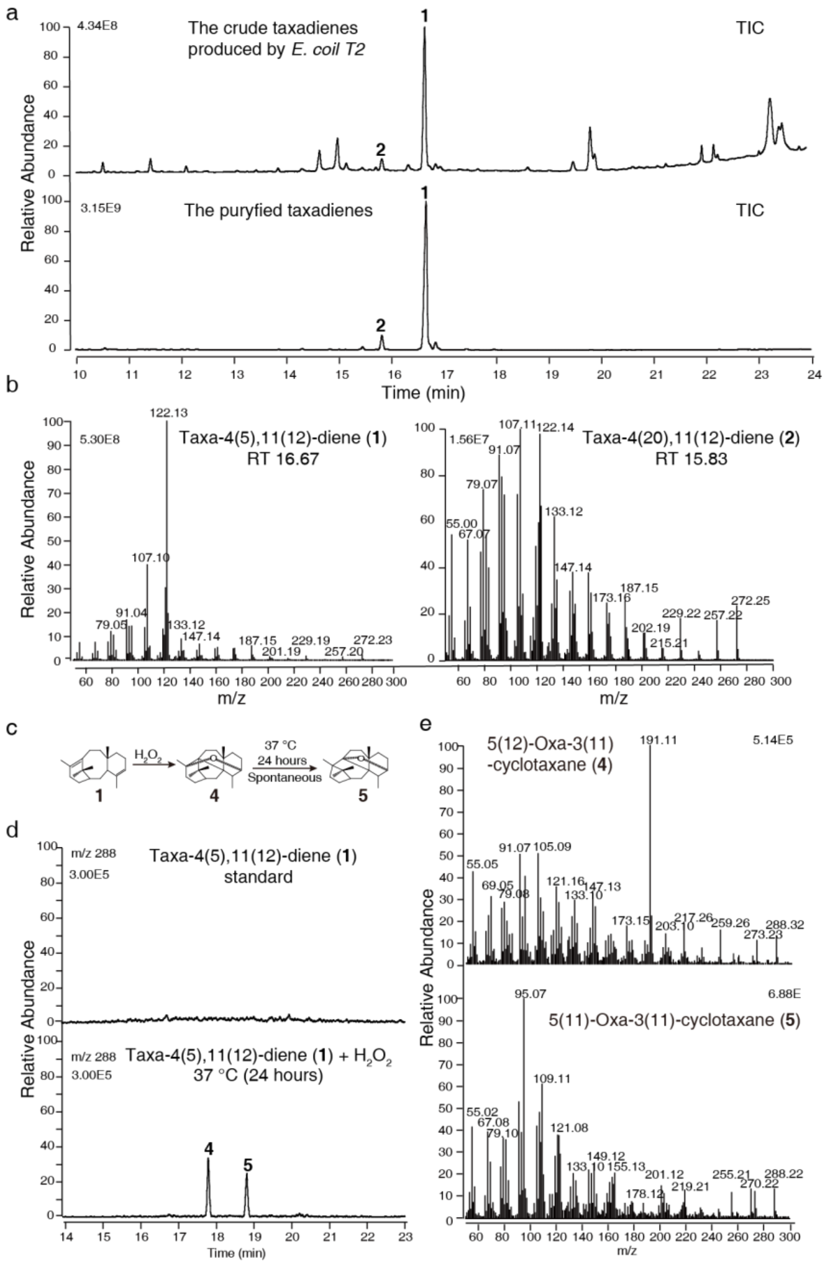


**Supplementary Fig. 25 Preparation of taxane intermediates.** (a) GC-MS analysis of the crude taxadiene produced by *E. coli* T2[25] and the purified taxadiene by thin-layer chromatography. Peaks **1** and **2** are taxa-4(5),11(12)-diene and taxa-4(20),11(12)-diene, respectively, produced by *E. coli* T2; (b) The MS/MS fragmentation of peaks **1** and **2**. (c) Schematic diagram of the oxidation reaction of taxadiene. OCT (**4**) was oxidized from **1** by hydrogen peroxide, and **4** spontaneously formed iso-OCT (**5**) at 37°C overnight. (d) GC-MS analysis of preparative **4** and **5**. Chromatograms at the extracted ions of $m/z^+$ 288. (e) The MS/MS fragmentation of peaks **4** and **5**.

### 3. **Supplementary Tables**

**Supplementary Table 1 The Taxus genome assembly.**

**A. Sequencing data used to assemble the Taxus genome.**

| Material | Data Type | Size (Gb) |
|---|---|---|
| Endosperm | IIIumina (WGS) | 693.73 |
| Endosperm | Pacbio (CLR) | 318.05 |
| Leaf | Pacbio (HiFi) | 26.06 |
| Endosperm | Hi-C | 1135.15 |
| Total | | 2172.99 |

WGS, Whole-Genome Sequencing; CLR, Continuous Long Read; HiFi, High fidelity reads.

**B. The statistics of the PacBio (CLR) assembly.**

| | Contig | Scaffold |
|---|---|---|
| Total | 10,232,190,857 bp | 10,237,952,243 bp |
| Count | 15,880 | 3,813 |
| Average | 644,344.51 bp | 2,685,012.39 bp |
| Max | 21,889,323 bp | 1,051,374,208 bp |
| N25 | 4,384,376 bp | 957,407,293 bp |
| N50 | 2,436,244 bp | 903,737,476 bp |
| N75 | 1,124,632 bp | 769,507,007 bp |
| N90 | 426,395 bp | 659,955,358 bp |
| N95 | 216,560 bp | 411,570,604 bp |

**Supplementary Table 2 Statistics of Hi-C sequencing and mapping.**

**A. Raw data aligned to the genome.**

| Sample | Hi-C data |
|---|---|
| Clean Paired-end Reads | 3,783,831,838 |
| Unmapped Paired-end Reads | 231,750,895 |
| Unmapped Paired-end Reads Rate (%) | 6.12 |
| Paired-end Reads with Singleton | 691,566,783 |
| Paired-end Reads with Singleton Rate (%) | 18.28 |
| Multi Mapped Paired-end Reads | 791,574,538 |
| Multi Mapped Paired-end Reads Ratio (%) | 20.92 |
| Unique Mapped Paired-end Reads | 2,068,939,622 |
| Unique Mapped Paired-end Reads Ratio (%) | 54.68 |

**B. Valid data of reads aligned to the genome.**

| Sample | Hi-C data |
|---|---|
| Unique Mapped Paired-end Reads | 2,068,939,622 |
| Dangling End Paired-end Reads | 47,562,609 |
| Dangling End Paired-end Reads Rate (%) | 2.30 |
| Self Circle Paired-end Reads | 5,581,545 |
| Self Circle Paired-end Reads Rate (%) | 0.27 |
| Dumped Paired-end Reads | 10,525,787 |
| Dumped Paired-end Reads Rate (%) | 0.51 |
| Interaction Paired-end Reads | 2,005,269,681 |
| Interaction Paired-end Reads Rate (%) | 96.92 |
| Valid Paired-end Reads | 2,001,492,791 |
| Valid Paired-end Reads Rate (%) | 96.74 |

**Supplementary Table 3 The lengths of pseudochromosomes generated in the Hi-C assembly.**

| Psudochromosome | Length (bp) |
|---|---|
| Chr01 | 1,052,277,571 |
| Chr02 | 995,865,175 |
| Chr03 | 958,316,394 |
| Chr04 | 950,298,798 |
| Chr05 | 904,642,436 |
| Chr06 | 909,132,773 |
| Chr07 | 786,775,274 |
| Chr08 | 774,176,834 |
| Chr09 | 770,396,028 |
| Chr10 | 681,618,719 |
| Chr11 | 660,890,931 |
| Chr12 | 412,525,553 |
| Total anchored length | 9,856,916,486 |
| Unanchored length | 381,035,757 |
| Total length | 10,237,952,243 |

**Supplementary Table 4 Assembly and annotation statistics of the Taxus genome.**

| Parameter | Value |
|---|---|
| Genome size | 10.23 Gb |
| GC content | 36.78% |
| Contig number | 15,879 |
| N50 length (contig) | 2.44 Mb |
| N90 length (contig) | 426.40 kb |
| Longest contig | 21,889,323 bp |
| Gene number | 42,746 |
| Average gene length | 13,260 bp |
| Chromosomes | 12 |
| Exons per gene | 3.5 |
| Total anchored | 9,856,916,486 bp |
| Repeat | 76.09% |
| Anchored rate | 96.28% |

**Supplementary Table 6 BUSCO analysis.**

**A. BUSCO analysis of genome annotation (embryophyte_odb10 (2020-9-10)).**

| | **Number** | **Rate (%)** |
|---|---|---|
| Complete BUSCOS (C) | 1,052 | 65.2 |
| Complete and single-copy BUSCOS (S) | 980 | 60.7 |
| Complete and duplicated BUSCOS (D) | 72 | 4.5 |
| Fragmented BUSCOs (F) | 265 | 16.4 |
| Missing BUSCOS (M) | 297 | 18.4 |
| Total BUSCO groups searched | 1,614 | 100.0 |

**B. BUSCO analysis of the gymnosperm species with genome data.**

| Species | Genome Length (Gb) | Contig N50 (kb) | C (%) | S (%) | D (%) | F (%) | M (%) |
|---|---|---|---|---|---|---|---|
| *Pinus lambertiana*[31] | 31.0 | 246.6 | 73.0 | 67.9 | 5.1 | 7.1 | 19.9 |
| *Pinus taeda*[32] | 20.6 | 25.4 | 40.7 | 35.9 | 4.8 | 18.6 | 40.7 |
| *Picea abies*[33] | 19.6 | 4.9 | 27.4 | 24.0 | 3.4 | 25.8 | 46.8 |
| *Abies alba*[34] | 18.16 | 14.1 | 15.4 | 12.5 | 2.9 | 16.9 | 67.7 |
| *Pseudotsuga menziesii*[35] | 16.6 | 44.1 | 67.8 | 62.5 | 5.3 | 11.0 | 21.2 |
| *Ginkgo biloba*[36] | 10.61 | 48.2 | 69.4 | 61.3 | 8.1 | 10.3 | 20.3 |
| *Taxus chinensis* var. *mairei* | 9.9 | 2435.93 | 65.2 | 60.7 | 4.5 | 16.4 | 18.4 |
| *Sequoiadendron giganteum*[37] | 8.13 | 347.95 | 50.7 | 47.5 | 3.2 | 14.9 | 34.4 |
| *Gnetum montanum*[38] | 4.07 | 25.02 | 83.0 | 79.3 | 3.7 | 4.1 | 12.9 |

Abbreviation: C, Complete BUSCOs; S, Complete and single-copy BUSCOs; D, Complete and duplicated BUSCOs; F, Fragmented BUSCOs; M, Missing BUSCOs.

**Supplementary Table 7 Functional annotation of Taxus genes.**

| Database | Number of annotated genes | Percentage (%) |
|---|---|---|
| GO | 13,694 | 32.04 |
| InterProScan | 33,210 | 77.69 |
| NR | 32,802 | 76.74 |
| Swiss-Port | 28,093 | 65.72 |
| TAIR | 31,126 | 72.82 |
| Total | 36,518 | 85.43 |

Abbreviation: GO, Gene Ontology Database; NR, Non-Redundant Proteins; Swiss-Port, Swiss-Port Database. TAIR, The Arabidopsis Information Resource.

**Supplementary Table 8 Statistics of repeated elements in the Taxus genome.**

| Classification | Number | Length (bp) | Percent of repeats (%) | Percent of genome (%) |
|---|---|---|---|---|
| Class I: Retroelement | 2,712,885 | 4,219,248,359 | 54.16 | 41.21 |
| LTR Retrotransposon | 2,571,938 | 4,080,706,163 | 52.38 | 39.86 |
| Ty1/Copia | 473,313 | 504,043,663 | 6.47 | 4.92 |
| Ty3/Gypsy | 2,082,710 | 3,561,506,787 | 45.72 | 34.79 |
| Other | 15,915 | 15,155,713 | 0.19 | 0.15 |
| Non-LTR Retrotransposon | 140,947 | 138,542,196 | 1.78 | 1.35 |
| LINE | 140,871 | 138,531,871 | 1.78 | 1.35 |
| SINE | 76 | 10,325 | 0.00 | 0.00 |
| Class II: DNA elements | 299,098 | 283,535,871 | 3.64 | 2.77 |
| CMC | 7,686 | 1,266,320 | 0.02 | 0.01 |
| hAT | 81,774 | 69,108,347 | 0.89 | 0.68 |
| PIF/Harbinger | 219 | 124,131 | 0.00 | 0.00 |
| MuLE-MuDR | 135,519 | 158,064,978 | 2.03 | 1.54 |
| Sola | 19,436 | 25,932,250 | 0.33 | 0.25 |
| Helitron | 36,248 | 26,622,352 | 0.34 | 0.26 |
| Enspm | 17,812 | 2,384,418 | 0.03 | 0.02 |
| Mariner | 404 | 33,075 | 0.00 | 0.00 |
| Small RNA | 502 | 73,190 | 0.00 | 0.00 |
| Simple repeats | 53,662 | 26,564,977 | 0.34 | 0.26 |
| Unknown | 5,208,975 | 2,838,802,940 | 0.34 | 0.26 |
| Total repeat fraction | 8,416,069 | 7,790,303,404 | 100.00 | 76.09 |

**Supplementary Table 27 Gene cluster identified through PlantiSMASH in the Taxus genome.**

| Type | Number | Chromosome |
|---|---|---|
| putative | 9 | 1, 3, 9, 10 |
| terpene | 7 | 1, 3, 6, 8, 9 |
| saccharide | 13 | 2, 6, 8, 9, 10, 12 |
| alkaloid | 1 | 1 |
| saccharide-polyketide | 1 | 2 |
| terpene-alkaloid | 1 | 4 |
| lignan-terpene | 1 | 5 |
| saccharide-terpene | 1 | 10 |

**4. Additional Data Supplementary Tables (separate file)**

**Supplementary Table 5 Genes expression dataset in Taxus.**

**Supplementary Table 9 Gene family analysis in 11 representative species.**

**Supplementary Table 10 Gene families gained in Taxus.**

**Supplementary Table 11 KEGG enrichment of the gained gene families.**

**Supplementary Table 12 Gene Ontology enrichment for significantly expanded gene families in Taxus.**

**Supplementary Table 13 The expanded gene families in the Taxus genome.**

**Supplementary Table 14 Pfam enrichment analysis of expanded gene families in Taxus.**

**Supplementary Table 15 KEGG enrichment analysis of expanded gene families in Taxus.**

**Supplementary Table 16 Classification of CYP450s in the Taxus genome.**

**Supplementary Table 17 CYP450 numbers in 69 representative plant genomes.**

**Supplementary Table 18 CYP450 gene groups in the Taxus.**

**Supplementary Table 19 Gene expression of CYP725As in groups 9.1 and 9.2.**

**Supplementary Table 20 Taxus gene clusters analyzed by PlantiSMASH.**

**Supplementary Table 21 Amino acid and nucleotide sequence alignments of TSs and T5αHs.**

**Supplementary Table 22 Genes screened by the gene-to-gene correlation analysis.**

**Supplementary Table 23 The known genes that are involved in paclitaxel biosynthesis.**

**Supplementary Table 24 Candidate genes potentially involved in paclitaxel metabolism.**

**Supplementary Table 25 RNA-seq data of tissues and mapping rates to the Taxus genome.**

**Supplementary Table 26 BAHD acyltransferase genes in the Taxus.**

**Supplementary Table 28 Primers used in the study.**

### 5. Supplementary References 1-38


1 Li, Y. *et al.* Induction of half-sib embryonic callus and production of taxiod compounds from *Taxus chinensis* var. *mairei*. *Int. J. Agric. Biol.* **21**, 719-725, doi:10.17957/IJAB/15.0949 (2019).

2 Marcais, G. & Kingsford, C. A fast, lock-free approach for efficient parallel counting of occurrences of k-mers. *Bioinformatics* **27**, 764-770, doi:10.1093/bioinformatics/btr011 (2011).

3 Li, H. Minimap2: pairwise alignment for nucleotide sequences. *Bioinformatics* **34**, 3094-3100, doi:10.1093/bioinformatics/bty191 (2018).

4 Danecek, P. *et al.* Twelve years of SAMtools and BCFtools. *Gigascience* **10**, 1-4, doi:10.1093/gigascience/giab008 (2021).

5 Xu, Z. & Wang, H. LTR_FINDER: an efficient tool for the prediction of full-length LTR retrotransposons. *Nucleic Acids Res.* **35**, 265-268, doi:10.1093/nar/gkm286 (2007).

6 Ellinghaus, D., Kurtz, S. & Willhoeft, U. LTRharvest, an efficient and flexible software for de novo detection of LTR retrotransposons. *BMC Bioinform.* **9**, 18, doi:10.1186/1471-2105-9-18 (2008).

7 Ou, S. & Jiang, N. LTR_retriever: a highly accurate and sensitive program for identification of long terminal repeat retrotransposons. *Plant Physiol.* **176**, 1410-1422, doi:10.1104/pp.17.01310 (2018).

8 Potter, S. C. *et al.* HMMER web server: 2018 update. *Nucleic Acids Res.* **46**, 200-204, doi:10.1093/nar/gky448 (2018).

9 Edgar, R. C. MUSCLE: multiple sequence alignment with high accuracy and high throughput. *Nucleic Acids Res.* **32**, 1792-1797, doi:10.1093/nar/gkh340 (2004).

10 Arkin, A. P. FastTree 2 - approximately maximum-likelihood trees for large alignments. *PLoS One* **5**, e9490, doi:10.1371/journal.pone.0009490 (2010).

11 Keller, I., Bensasson, D. & Nichols, R. A. Transition-transversion bias is not universal: a counter example from grasshopper pseudogenes. *PLoS Genet.* **3**, e22, doi:10.1371/journal.pgen.0030022 (2007).

12 Rice, J., Longden, I. & Bleasby, A. EMBOSS: the European molecular biology open software suite. *Trends Genet.* **16**, 276-277, doi: 10.1016/s0168-9525(00)02024-2 (2000).

13 De La Torre, A. R., Li, Z., Van de Peer, Y. & Ingvarsson, P. K. Contrasting rates of molecular evolution and patterns of selection among gymnosperms and flowering plants. *Mol. Biol. Evol.*, 1363-1377, doi:10.1093/molbev/msx069 (2017).

14 Kim, D., Langmead, B. & Salzberg, S. L. HISAT: a fast spliced aligner with low memory requirements. *Nat. Methods* **12**, 357-360, doi:10.1038/nmeth.3317 (2015).

15 Pertea, M., Kim, D., Pertea, G. M., Leek, J. T. & Salzberg, S. L. Transcript-level expression analysis of RNA-seq experiments with HISAT, StringTie and Ballgown. *Nat. Protoc.* **11**, 1650-1667, doi:10.1038/nprot.2016.095 (2016).

16 Robinson, M. D., McCarthy, D. J. & Smyth, G. K. edgeR: a Bioconductor package for differential expression analysis of digital gene expression data. *Bioinformatics* **26**, 139-140, doi:10.1093/bioinformatics/btp616 (2009).

17 Wegrzyn, J. L., Lee, J. M., Tearse, B. R. & Neale, D. B. TreeGenes: a forest tree genome database. *Int. J. Plant Genomics* **2008**, 412875, doi:10.1155/2008/412875 (2008).

18 Altschul, S. F., Gish, W., Miller, W., Myers, E. W. & Lipman, D. J. Basic local alignment search tool. *J. Mol. Biol.* **215**, 403-410, doi:10.1016/S0022-2836(05)80360-2 (1990).

19 Li, L., Stoeckert, C. J., Jr. & Roos, D. S. OrthoMCL: identification of ortholog groups for eukaryotic genomes. *Genome Res.* **13**, 2178-2189, doi:10.1101/gr.1224503 (2003).

20 Edgar, R. MUSCLE: multiple sequence alignment with high accuracy and high throughput. *Nucleic Acids. Res.* **32**, 1792-1797, doi:10.1093/nar/gkh340 (2004).

21 Castresana, J. Selection of conserved blocks from multiple alignments for their use in phylogenetic analysis. *Mol. Biol. Evol.* **17**, 540-552, doi:10.1093/oxfordjournals.molbev.a026334 (2000).

22 Stamatakis, A. RAxML version 8: a tool for phylogenetic analysis and post-analysis of large phylogenies. *Bioinformatics* **30**, 1312-1313, doi:10.1093/bioinformatics/btu033 (2014).

23 Sanderson, M. J. r8s: inferring absolute rates of molecular evolution and divergence times in the absence of a molecular clock. *Bioinformatics* **19**, 301-302, doi:10.1093/bioinformatics/19.2.301 (2003).

24 De Bie, T., Cristianini, N., Demuth, J. P. & Hahn, M. W. CAFE: a computational tool for the study of gene family evolution. *Bioinformatics* **22**, 1269-1271, doi:10.1093/bioinformatics/btl097 (2006).

25 Yu, G., Wang, L. G., Han, Y. & He, Q. Y. clusterProfiler: an R package for comparing biological themes among gene clusters. *OMICS* **16**, 284-287, doi:10.1089/omi.2011.0118 (2012).

26 Urban, P., Mignotte, C., Kazmaier, M., Delorme, F. & Pompon, D. Cloning, yeast expression, and characterization of the coupling of two distantly related *Arabidopsis thaliana* NADPH-Cytochrome P450 reductases with P450 CYP73A5. *J. Biol. Chem.* **272**, 19176-19186, doi:10.1074/jbc.272.31.19176 (1997).

27 Pompon, D., Louerat, B., Bronine, A. & Urban, P. in *Methods in Enzymology* Vol. **272** (eds Eric F. Johnson & Michael R. Waterman) 51-64 (Academic Press, 1996).

28 Biggs, B. W. *et al.* Orthogonal assays clarify the oxidative biochemistry of taxol P450 CYP725A4. *ACS Chem. Biol.* **11**, 1445-1451, doi:10.1021/acschembio.5b00968 (2016).

29 Bian, G. *et al.* Production of taxadiene by engineering of mevalonate pathway in *Escherichia coli* and endophytic fungus *Alternaria alternata* TPF6. *Biotechnol. J.* **12**, doi:10.1002/biot.201600697 (2017).

30 Kuang, X., Sun, S., Wei, J., Li, Y. & Sun, C. Iso-Seq analysis of the *Taxus cuspidata* transcriptome reveals the complexity of Taxol biosynthesis. *BMC Plant Biol.* **19**, 210, doi:10.1186/s12870-019-1809-8 (2019).

36 Guan, R. et al. Draft genome of the living fossil *Ginkgo biloba*. *Gigascience* **5**, 49, doi:10.1186/s13742-016-0154-1 (2016).

37 Scott, A. D. et al. A reference genome sequence for giant sequoia. *G3* **10**, 3907-3919, doi:10.1534/g3.120.401612 (2020).

38 Wan, T. et al. A genome for gnetophytes and early evolution of seed plants. *Nat. Plants* **4**, 82-89, doi:10.1038/s41477-017-0097-2 (2018).
